# Low parental conflict, no endosperm hybrid barriers, and maternal bias in genomic imprinting in selfing *Draba* species

**DOI:** 10.1101/2024.01.08.574548

**Authors:** Renate M. Alling, Katrine N. Bjerkan, Jonathan Bramsiepe, Michael D. Nowak, A. Lovisa S. Gustafsson, Christian Brochmann, Anne K. Brysting, Paul E. Grini

## Abstract

In flowering plants, a distinct post-zygotic hybridization barrier between closely related species can arise during seed maturation, resulting in embryo lethality due to abnormal endosperm development. The endosperm initially works as a nutrient sink, acquiring nutrients from adjacent tissues, but later undergoes cellularization, switching to serve as a nutrient source. In hybrid seeds, this cellularization switch can be hampered if the endosperm genomic ratio is imbalanced. Disruption in the genomic ratio can be caused when species of different ploidy are crossed, but also by crosses between species with identical ploidy, if the effective ploidy differs. One factor proposed to influence effective ploidy is the epigenetic phenomenon genomic imprinting, the parent-of-origin specific expression of alleles inherited either maternally or paternally. It has been proposed that outbreeding species exhibit higher effective ploidy compared to selfing species, as a consequence of parental conflict in resource allocation to the developing progenies. This suggests a low anticipation of endosperm-based post-zygotic hybridization barriers between selfing species of similar ploidy. Here, we show that in crosses between the diploid selfing arctic species *Draba fladnizensis*, *D. nivalis* and *D. subcapitata*, the endosperm-based post-zygotic hybridization barrier is absent, supporting low parental conflict. To investigate parent-of-origin allele specific expression, we conducted a genomic imprinting study in *D. nivalis* and compared to previous studies in other Brassicaceae species. We report a high number of maternally expressed genes (MEGs) and concomitantly low numbers of paternally expressed genes (PEGs). Our results suggest rapid evolution of MEGs and loss of PEGs in a mating system with low parental conflict, proposing that selfing arctic species may exhibit a generally stronger maternal expression bias as an adaptive mechanism to efficiently cope with an extreme environment.

## Introduction

Speciation is a continuous process towards reproductive isolation between two lineages, obtained by the evolution of hybridization barriers prior to (pre-zygotic) or following fertilization (post-zygotic) (Rieseberg & Willis, 2007; Florez-Rueda *et al.*, 2016). In flowering plants, a distinct post-zygotic hybridization barrier between closely related species can arise during seed maturation (Lafon-Placette & Köhler, 2014), resulting in embryo lethality due to abnormal endosperm development (Cooper & Brink, 1942). This endosperm-based hybridization barrier is found in several genera across the angiosperm clade, including *Oryza* (Ishikawa *et al.*, 2011; Sekine *et al.*, 2013; Zhang *et al.*, 2016b; Tonosaki *et al.*, 2018; Wang *et al.*, 2018), *Arabidopsis* (Lafon-Placette & Köhler, 2016), *Capsella* (Rebernig *et al.*, 2015; Dziasek *et al.*, 2021), *Arabis* (Petrén *et al.*, 2023), *Solanum* (Johnston & Hanneman, 1982; Cornejo *et al.*, 2012; Florez-Rueda *et al.*, 2016; Roth *et al.*, 2019), and *Mimulus* (Oneal *et al.*, 2016; Flores-Vergara *et al.*, 2020; Kinser *et al.*, 2021), suggesting that it is controlled by a conserved mechanism.

The endosperm is a triploid tissue, composed of two maternal genomic copies for every paternal copy. During the initial stages of development, it forms a syncytium which acquires nutrients from adjacent tissues. After sufficient acquisition of nutrients, the endosperm nuclei are distributed into separate cells and when this cellularization process is finished, the endosperm works as a nutrient source for the embryo (Lafon-Placette & Köhler, 2014). Disruption of the genomic ratio of the endosperm, causing either a maternal or paternal excess, can affect the time point of cellularization and is believed to be the indirect cause for embryo lethality in interploidy crosses (Lin, 1984; Birchler, 1993; Scott *et al.*, 1998; Leblanc *et al.*, 2002). However, it has also been reported cases where interspecific crosses involving species of identical ploidy, i.e. homoploid crosses, have displayed features of maternal or paternal excess (Müntzing, 1933; Cooper & Brink, 1945; Marks, 1966; Williams & White, 1976; Sukno *et al.*, 1999). To explain how parental excess in homoploid interspecific crosses with identical ploidy can occur, (Johnston *et al.*, 1980) coined the term Endosperm Balance Number (EBN), which explains how the effective ploidy may differ from the expected ploidy. One factor proposed to influence effective ploidy, is the epigenetic phenomenon called imprinting (Haig & Westoby, 1991; Bushell *et al.*, 2003).

Genomic imprinting is parent-of-origin specific expression of alleles inherited either maternally or paternally (Hornslien *et al.*, 2019; Batista & Köhler, 2020). This epigenetic phenomenon is widespread in the angiosperm clade, including genera such as *Arabidopsis* (Wolff *et al.*, 2011; Pignatta *et al.*, 2014; Klosinska *et al.*, 2016; Del Toro-De León & Köhler, 2019; Hornslien *et al.*, 2019; Picard *et al.*, 2021; van Ekelenburg *et al.*, 2023), *Brassica* (Yoshida *et al.*, 2018; Liu *et al.*, 2018; Rong *et al.*, 2021), *Capsella* (Hatorangan *et al.*, 2016; Lafon-Placette *et al.*, 2018), *Fragaria* (Liu *et al.*, 2021), *Mimulus* (Xu *et al.*, 2014; Kinser *et al.*, 2021), *Oryza* (Zhang *et al.*, 2016b; Wang *et al.*, 2018), *Ricinus* (Xu *et al.*, 2014), *Solanum* (Florez-Rueda *et al.*, 2016; Roth *et al.*, 2019)*, Sorghum* (Zhang *et al.*, 2016a) and *Zea* (Zhang *et al.*, 2014; Dong *et al.*, 2023).

One theory on the evolution of genomic imprinting in the endosperm is the “kinship theory” (Haig & Westoby, 1991), which is based on a difference of interest in resource allocation to the embryo between the paternal and maternal parents. Parental conflict is more pronounced in outbreeding species, in which the female may be fertilized by several males, and it is thus not relevant for strongly inbreeding (selfing) species. Notably, the transition from outbreeding to selfing is a common evolutionary change in plants (Barrett, 2002). The weak inbreeder/strong outbreeder (WISO) hypothesis is based on differences in parental conflict between outbreeding and selfing populations or species (Brandvain & Haig, 2005, 2018; Raunsgard *et al.*, 2018), and is supported by recent evidence. In *Arabis alpina*, crosses between selfing and outbreeding populations showed an endosperm-based hybridization barrier, whereas crosses between populations with the same mating system had significantly higher success rate (Petrén *et al.*, 2023). In *Capsella*, interspecific hybrids between obligate outbreeders, recent selfers, and ancient selfers, the parental conflict was observed as precocious or delayed cellularization of the endosperm, and decreased with a longer evolutionary history of selfing (Lafon-Placette *et al.*, 2018). In intraspecific hybrids between *Dalechampia scandens* populations with different outbreeding intensity, seed size was positively correlated with the paternal seed size from more outbreeding populations when they were crossed with maternal individuals from less outbreeding populations (Raunsgard *et al.*, 2018). A parental conflict in interspecific crosses between a maternal selfing species and a paternal outbreeder, can be explained by an increase in paternally expressed genes (PEGs) causing paternal excess. Both outbreeding species from *Capsella* and *Arabidopsis* have shown an increase in PEGs when compared to selfing species from the same genus. However, there is no difference in maternally expressed genes (MEGs) in species with different mating systems, and therefore, it cannot explain how a maternal outbreeder can cause a maternal excess when crossed with a paternal selfer (Klosinska *et al.*, 2016; Lafon-Placette *et al.*, 2018).

Studies investigating genomic imprinting and post-zygotic barriers as a result of parental conflict have predominantly been conducted in the large Brassicaceae family. This family has a global distribution excluding Antarctica (Lysak & Koch, 2011) and encompasses numerous economically important species with diverse applications in medicine, agriculture, and scientific research (Raza *et al.*, 2020). A recent phylogeny of Brassicaceae (Hendriks *et al.*, 2023) divides the family into five supertribes: Camelinodae, Brassicodae, Hesperodae, Arabodae and Heliophilodae (Figure 1 A). Research on the endosperm-based hybridization barrier has focused on the supertribes Camelinodae and Arabodae, whereas imprinting studies have been conducted in Camelinodae and Brassicodae.

**Figure 1:**
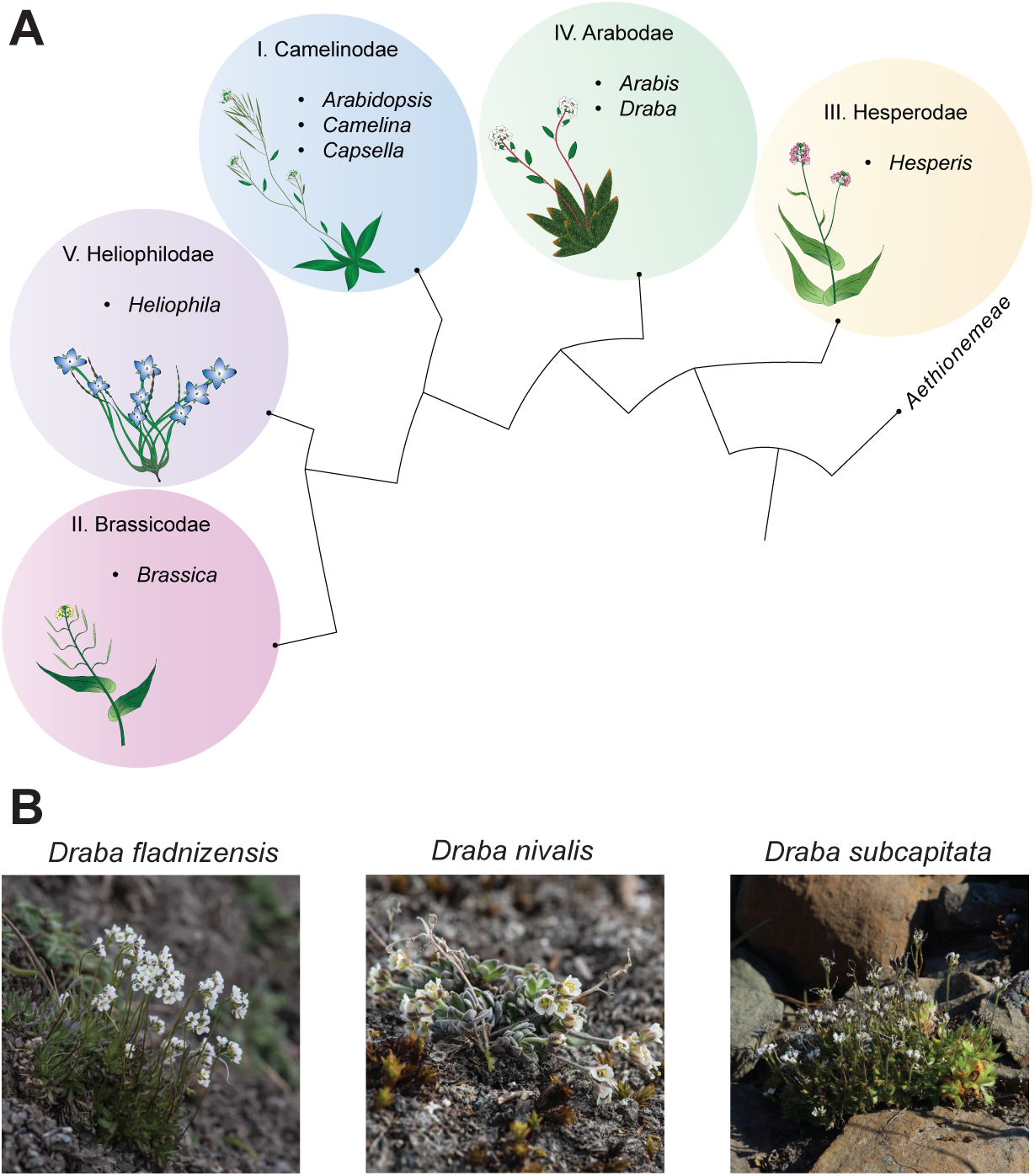
Phylogeny of the Brassicaceae family and photos of the target species of this study, *Draba fladnizensis*, *D. nivalis* and *D. subcapitata*. **(A)** Nuclear Brassicaceae phylogeny adapted from (Hendriks *et al.*, 2023), dividing the family into five supertribes (I-IV), with Aethionemeae as outgroup. Within each supertribe, representative and important genera are listed and illustrated. (B) *Draba fladnizensis*, *D. nivalis* and *D. subcapitata*, photographed by Geir Arnesen and retrieved from (Elven *et al.*, 2020).

Within supertribe Arabodae, we find *Draba* which is the largest Brassicaceae genus (Figure 1 A). It encompasses approximately 370 species predominantly from alpine and arctic regions (Warwick *et al.*, 2006), and shows a high rate of polyploidization and speciation, hypothesized to have been driven by the frequent glacial cycles during the Pleistocene (Al-Shehbaz *et al.*, 2006; Bailey *et al.*, 2006; Jordon-Thaden & Koch, 2008). Selfing is beneficial in harsh environments with limited or no pollinators and is widespread in arctic plants (Solbrig, 1980). Notable examples are the circumpolar species *Draba nivalis*, *D. subcapitata* and *D. fladnizensis* (Figure 1 B), which are diploid with eight chromosome pairs and predominantly selfing (Brochmann, 1993; Brochmann *et al.*, 1993). Many recently evolved cryptic species isolated by F_1_ sterility barriers have been identified within each of these taxonomically well-defined species of *Draba*, whose selfing mating system is likely to have induced high rates of accumulation of hybrid incompatibilities (Brochmann *et al.*, 1993; Grundt *et al.*, 2006; Marie-Orleach *et al.*, 2022, 2023). Quantitative trait locus (QTL) mapping in *Draba nivalis* revealed a complex genetic architecture of such post-zygotic reproductive isolation, including underdominant loci most likely due to microchromosomal rearrangements as well as nuclear–nuclear and cyto–nuclear epistatic interactions between loci, indicating Bateson–Dobzhansky–Muller (BDM) incompatibilities (Skrede *et al.*, 2008; Gustafsson *et al.*, 2014). Thus, one might expect such incipient species in the Arctic to be deeply divergent, but on the contrary, (Gustafsson *et al.*, 2022) estimated that some of them may have diverged even during the last few millennia. Not surprisingly, in addition to strong F_1_ hybrid sterility barriers within each of these three species of *Draba*, there are also strong F_1_ hybrid sterility barriers between them (Mulligan, 1974), which probably are of recent Pleistocene origin (Grundt *et al.*, 2004, 2006). There is also some evidence suggesting pre-zygotic or post-zygotic barriers between the three species affecting seed viability. In the study by (Brochmann *et al.*, 1993), crosses between them did either not result in any fertilized ovules or in only a few viable seeds. It is thus possible that also an endosperm-based hybridization barrier might be present between these species.

With a new genome resource for *D. nivalis* in place (Nowak *et al.*, 2020), previously reported hybrid barriers, high speciation rates, and a selfing mating system, these three *Draba* species represent an interesting system to study endosperm-based hybridization barriers and parental conflict in relation to genomic imprinting. The species are likely to experience little parental conflict because of high levels of self-fertilization, and to test this we analyzed endosperm-based hybridization barriers in Draba interspecific crosses To investigate parent-of-origin allele specific expression, we conducted a genomic imprinting study in *D. nivalis* and compared to previous studies in other Brassicaceae species. We found however no significant endosperm-based species barrier in our interspecific hybrid seeds, but rather a high number of maternally expressed genes (MEGs) and concomitantly low numbers of PEGs. These results show rapid evolution of MEGs and loss of PEGs in a mating system with low parental conflict, suggesting that selfing arctic species may exhibit a generally stronger maternal expression bias as an adaptive mechanism to efficiently cope with an extreme environment.

## Materials and Methods

### Plant material, seed sterilization and growth conditions

Information about the geographic origin of the *Draba* populations used for hybrid crosses and imprinting analysis can be found in Figure 2 and Supplementary Table S1. Seed sterilization and scarification were performed as established by (AU-Lindsey *et al.*, 2017), except that the scarification process with bleach was prolonged from 5 to 30 min. This step was important to increase germination rate. After sterilization, seeds were transferred to petri-dishes containing ½ MS growth medium supplemented with 2% sucrose (Murashige & Skoog, 1962). Afterwards the seeds underwent a 2-day stratification period at 4°C, before germination at 22°C with a 16h/8h light/dark cycle. After two weeks, seedlings were transferred to soil containing approximately 30% sand and cultivated at 15°C (16h/8h light/dark cycle, 160 µmol/m2/sec, 60-65% humidity). When the plants had developed a healthy rosette, approximately 3-4 weeks after transfer to soil, they underwent a two-month vernalization period, following the approach described by (Skrede *et al.*, 2008). Before crossing the plants, they were placed in a growth chamber at 15°C (16h/8h light/dark cycle, 160 µmol/m2/sec, 60-65% humidity) for 3-4 weeks until flower bud production and kept under the same conditions during silique maturation.

**Figure 2:**
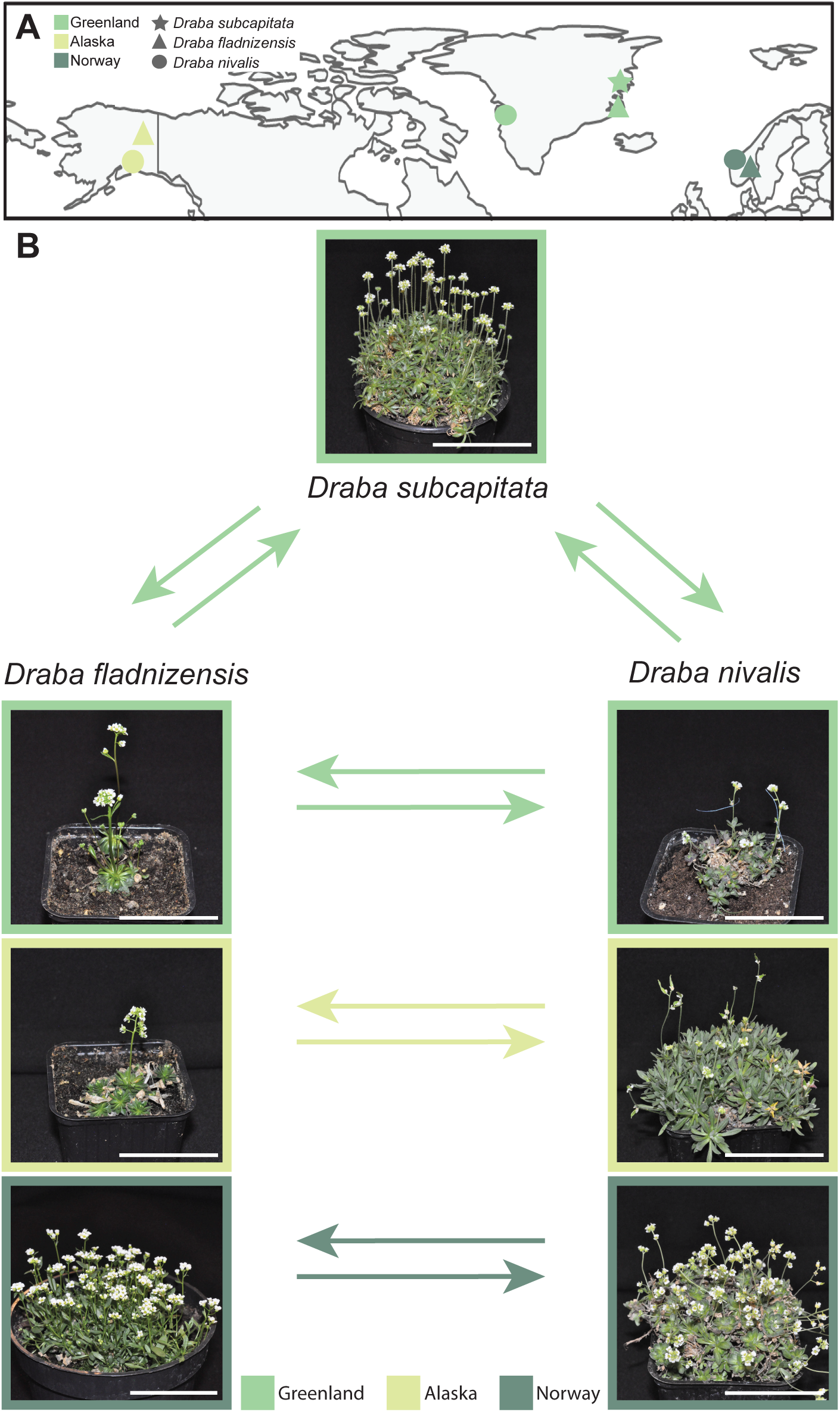
Geographic locations and crossing scheme of the three *Draba* species used in this study. **(A)** Sampling locations of *D. subcapitata* (star), *D. fladnizensis* (triangle) and *D. nivalis* (circle) from Greenland (light green), Alaska (yellow) and Norway (dark green). See Supplementary Table S1 for details. **(B)** Crossing scheme with photos of representative individuals used for interspecific crosses within Greenland, Alaska and Norway populations. Cross direction is indicated with arrows. Scale bar = 5 cm.

### Crossing and germination assays

Intra- and interspecific crossing experiments were performed according to the methodology established by (Grundt *et al.*, 2006). Emasculation was performed before the buds started opening to avoid self-fertilization, and hand-pollinated after two days. For each cross combination, we used 4-16 different individual plants to produce approximately 100 seeds, and each silique was harvested individually. For the germination assay, seeds from each silique were counted and placed on a petri-dish with ½ MS growth-medium and scored for germination after 14 days at 22°C. For the imprinting study, four individuals of each population were used for intra- and interspecific crosses aimed approximately 20 siliques for each reciprocal direction. Seeds were harvested at 7 days after pollination (DAP) and flash frozen in liquid nitrogen.

### RNA isolation, sequencing and GFP marker line generation

RNA was isolated from whole seeds as described in the RNeasy Plant Mini Kit (Qiagen). Sequencing libraries were prepared with TruSeq^TM^ RNA library prep kit (Illumina) and sequenced with Illumina HiSeq 4000, paired end (150 bp) with three lanes in total. The sequencing and library preparation were performed at the Norwegian Sequencing Center (NSC; https://www.sequencing.uio.no/). The Total Endosperm (*TE1*; AT4G00220; *proTE1>>H2A-GFP*) marker line (van Ekelenburg *et al.*, 2023) was transformed into a Norwegian *D. nivalis* population (Grimsdalen; Supplementary Table S1) using the floral dip method (Clough & Bent, 1998).

### Microscopy

To determine the developmental stage, seeds were cleared using a chloral hydrate:water:glycerol 8:2:1(w/v/v; (Grini *et al.*, 2002) solution and imaged using a Axioplan2 microscope. Mature dry seeds and siliques were imaged with a Leica Z16apoA microscope connected to a Nikon D90 camera. The *TE1-GFP* lines were imaged using the Andor DragonFly spinning disc confocal microscope with a Zyla4.2 sCMOS 2048x2048 camera attached with excitation of 488 nm and emission of 525 nm.

### Statistics and imprinting analysis

Area and mean area (mm^2^) of mature siliques and seeds (Supplementary Data S1) were measured and calculated by converting images to black and white and then using the ImageJ “Threshold” and “Analyze particles” functions (https://imagej.nih.gov/ij/). The resulting mean values, together with mean number of seeds per silique, were plotted as boxplots using ggplot2 (Wickham, 2016) and dplyr packages (Wickham *et al.*, 2023) in R-studio (version 2023.03.1 + 446; (R Core Team, 2023; RStudio Team, 2023). For the germination assay results, significance was calculated using Welch’s t-test (Welch, 1947). Midparent values were calculated from both parental datasets and the resulting median was indicated on the germination rate boxplots. Orthogroups between *D. nivalis* and *A. thaliana* were identified using OrthoFinder v2.5.2 tutorials (Emms & Kelly, 2019) (Supplementary Data S4). The map was generated using R version 4.3.2 (2023-10-31) together with the built-in tmap package version ‘3.3.3’ (R Core Team, 2023).

A detailed description of the imprinting analyses is provided as supplementary methods and a GitHub repository, available at https://github.com/PaulGrini/Draba. In summary, parental reads from each *D. nivalis* population (Alaska and Norway) were mapped to a *D. nivalis* Alaska transcriptome reference (Nowak *et al.*, 2020) and furthermore polished using pilon (Walker *et al.*, 2014), in order to generate population specific references. The population specific sequences were combined to one target file, to which the hybrid reads were mapped. Afterwards, the Informative Reads Pipeline (IRP) was utilized to identify and extract informative reads (Hornslien *et al.*, 2019).

The informative reads (Supplementary Data S5) were analyzed in R-studio (version 2023.03.1 + 446; (R Core Team, 2023) performing differential expression analysis using the R package limma (Ritchie *et al.*, 2015) and normalization factors generated with the IRP (Hornslien *et al.*, 2019). Venn diagrams and histograms were generated using VennDiagram version 1.7.3 (Chen & Boutros, 2011), and the hist() function in base R. DESeq2 (Love *et al.*, 2014) was used for differential expression analysis of the parental samples, using count reads generated in the IRP, to identify population-specific expression bias. A general seed coat (GSC) filter was applied before differential expression analysis in the IRP. This filter was based on a 4-fold GSC specific enrichment of transcripts in the seed of *A. thaliana* (Hornslien *et al.*, 2019). In addition, a DESeq2 analysis was performed on all samples, both heterozygous and homozygous, using count reads generated in the IRP, to generate a principal component analysis (PCA) plot using PCATools version 2.14.0 (Blighe *et al.*, 2019).

Identified imprinted genes (Supplementary Data S5) were compared to imprinted genes identified in other studies using Complex UpSetR version 1.3.3 (Krassowski *et al.*, 2022). A gene ontology (GO) analysis was performed on the overlapping MEGs identified between *D. nivalis* and any of the other studies (Supplementary Data S5), using ClusterProfiler version 4.8.2 (Wu *et al.*, 2021) and enrichplot version 1.20.0 (Yu, 2023) with standard settings.

### Accession Numbers and Supplemental data

All sequences generated in this study have been deposited to the National Center for Biotechnology Information Sequence Read Archive (https://www.ncbi.nlm.nih.gov/sra) with project number PRJNAxx. Supplemental data files and scripts are available from the Github depository (https://github.com/PaulGrini/Draba).

## Results

### Interspecific *Draba* crosses display no endosperm-based hybridization barrier

We investigated parental conflict in hybrid seeds between strong selfing *Draba* species. *D. nivalis*, *D. fladnizensis* and *D. subcapitata* populations from Alaska, Greenland and Norway (Figure 2 A) (Brochmann, 1993; Grundt *et al.*, 2006) (Supplementary Table S1) were selected or collected, and maintained at controlled environment conditions. To control for defects in fertility or fecundity in the populations at controlled growth conditions, we first inspected reproduction parameters in selfed populations. We found large variation in silique size both between species and between populations (Supplementary Figure 1 A), but the seed set was high (88%-100%; Supplementary Figure 1 B, Supplementary Data S1). The number of seeds per silique was characteristic for each species (Supplementary Figure 1 B) and in line with published data, albeit with a slightly higher number of seeds than what is reported from natural populations (Elven *et al.*, 2020). The seed size was inversely correlated with the number of seeds, with *D. subcapitata* having the largest seeds and smallest number of seeds (Supplementary Figure 1 C).

We reciprocally crossed all three species from Greenland, and *D. fladnizensis* and *D. nivalis* from both Alaska and Norway (Figure 2 B). We harvested separately 3-23 silicles individually from each cross. The seed set ranged from 66% to 97%, suggesting a weak pre-zygotic barrier in the reciprocal hybrids between both the Greenland and Alaska populations of *D. nivalis* and *D. fladnizensis* (Supplementary Data S1). The mature hybrid seeds varied in seed number and size, depending on parental species and the direction of the cross (Supplementary Figure 2). In reciprocal hybrids between the Greenland populations of *D. subcapitata* and *D. nivalis* or *D. fladnizensis*, the seeds were of the same size as those of the maternal parent. However, in all three populations investigated, the seeds of the reciprocal hybrids between *D. nivalis* and *D. fladnizensis* were either intermediate between or smaller than those of the parental species, or similar to those of the paternal parent (Supplementary Figure 1, Supplementary Figure 2).

We scored germination rate of seeds from individual siliques from each cross, using seeds obtained after spontaneous selfing of the parental plants as controls (Figure 3, Supplementary Figure 3). The germination rate of the seeds from the reciprocal crosses between *D. nivalis* and *D. fladnizensis* were not significantly different from those of the maternal parent, but significantly different from those of the paternal parent. However, no hybrid seed germination rates were significantly different from the midparent value, indicating an overall lack of a seed-based post-zygotic barrier. In the reciprocal crosses between *D. fladnizensis* and *D. subcapitata,* the only significant difference in germination rate was found between the paternal parent *D. subcapitata* and *D. fladnizensis* × *subcapitata.* Interestingly, the *D. nivalis* × *subcapitata* seeds had almost complete overall germination rate, significantly higher than both the maternal and paternal parent and the midparent value. Thus, no hybrid cross combinations showed presence of an endosperm-based hybridization barrier. We observed some significant differences from the paternal parent, but none from the maternal parent or midparent values.

**Figure 3:**
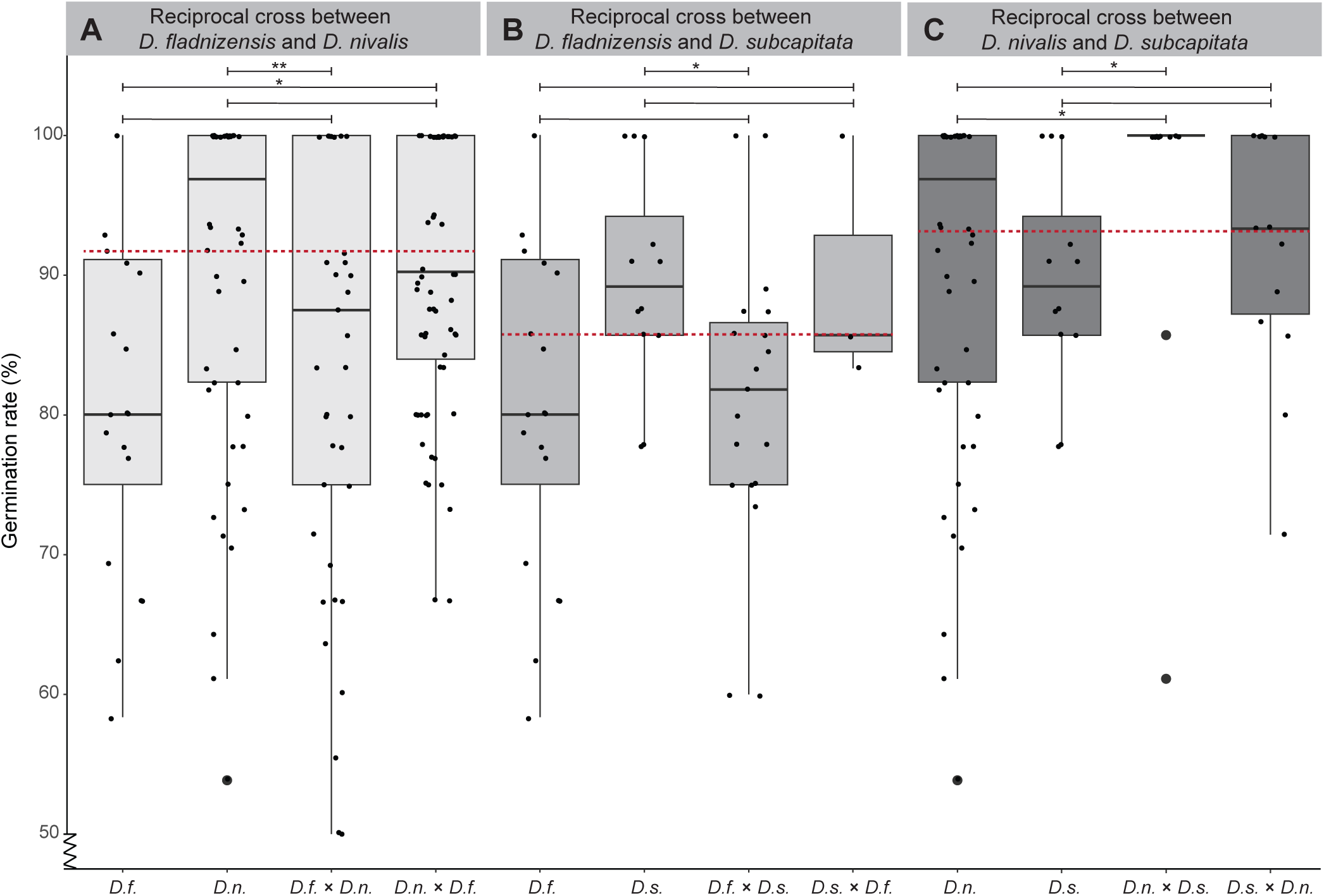
Germination rates of *Draba* hybrid seeds do not indicate endosperm-based hybridization barriers. Crosses are indicated as maternal × paternal below the x-axis. If the paternal cross partner is not indicated, the maternal and paternal individuals are the same. Biological replicates (n) are given by germination frequency per silique. **(A)** Germination rates of *D. fladnizensis* (*D.f.*), n = 20, and *D. nivalis* (*D.n.*), n = 50, and their reciprocal hybrids *D.f.* × *D.n.*, n = 41, and *D.n.* × *D.f.*, n = 68. **(B)** Germination rates of *D.f.*, n = 20, and *D. subcapitata* (*D.s.*), n = 12, and their reciprocal hybrids *D.f.* × *D.s.*, n = 19, and *D.s.* × *D.f.*, n = 3. **(C)** Germination rates of *D.n.*, n = 50, and *D.s*., n = 12, and their reciprocal hybrids *D.n.* × *D.s.*, n = 17, and *D.s.* × *D.n.*, n = 14. Biological replicates are plotted as small dots. Outliers are plotted as large dots. Midparent values are indicated by red dashed lines. Boxplots of the parental controls (*D.f., D.n., and D.s.)* are repeated for easier comparison and indication of significance tests. For comparison of germination rates within each region, see Supplementary Figure 3. Note that the Y-axis is truncated up to <50%. Significance is indicated for comparisons between reciprocal hybrids and parental lines (Welch’s t-test: *P ≤ 0.05; **P ≤ 0.01; not significant is indicated by no asterisk).

### *Draba nivalis* endosperm cellularization can be staged using a transgenic reporter

To investigate parental conflict and genomic imprinting in *Draba* species, we selected two populations of *D. nivalis* from Alaska and Norway. Recent assembly of the *D. nivalis* Alaska genome (Nowak *et al.*, 2020) allows detection of parent-of-origin allele-specific expression from intraspecies Alaska Norway hybrid seeds and *D. nivalis* was therefore the most suited *Draba* species for this analysis. To rule out the possibility of an incomplete pre-zygotic or seed-based post-zygotic hybridization barrier that could affect the genomic imprinting experiment, we investigated the seed phenotypic parameters of the intraspecific hybrid cross. The seed set in both parental populations and their reciprocal hybrids was near complete, indicating a lack of hybridization barriers (Supplementary Figure 4). The Alaska population had slightly larger seeds than the Norway population and the reciprocal hybrids followed the maternal parent in size, and the seed size was inversely correlated with seed number (Supplementary Figure 4).

For staging purposes, we examined seed development at seven and 12 DAP in the parental populations and the hybrids. They all showed a similar developmental progress, with primarily globular embryo stages and uncellularized endosperm present at seven DAP, and late heart embryo stages, presumably with cellularized endosperm, at 12 DAP (Supplementary Figure 5). We wanted to perform imprinting analysis before the onset of endosperm cellularization and therefore inspected seven DAP *D. nivalis* Alaska Norway hybrid seeds in detail. A similar developmental progression between the two reciprocal intraspecific hybrids was observed, with the majority of the seeds having an embryo at octant to globular stage and a primarily uncellularized endosperm (Figure 4). Both embryo and endosperm staging and nuclei counts suggested high similarity between the reciprocal intraspecific hybrids.

**Figure 4:**
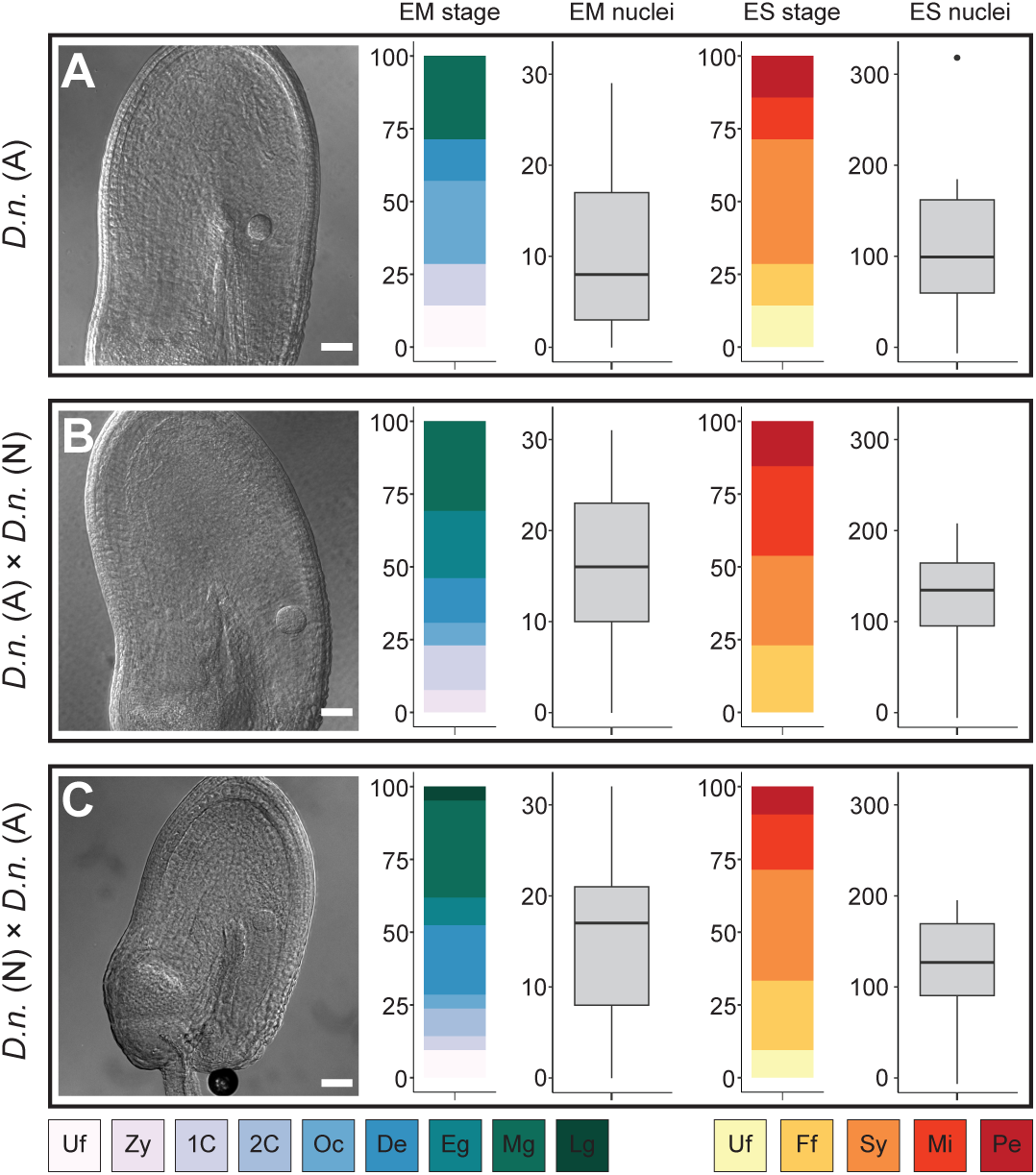
Embryo and endosperm stages and number of nuclei in *Draba nivalis* Alaska and reciprocal intraspecific hybrids between Alaska and Norway. **Each panel shows** (from left to right): light micrographs of seeds at seven days after pollination, relative frequencies of embryo (EM) stages with number of embryo nuclei and relative frequencies of endosperm (ES) stages and number of nuclei in the endosperm. Crosses are indicated as maternal × paternal. **(A)** *D. nivalis* Alaska (*D.n.*(A)) self cross, n = 7; **(B)** *D.n.*(A) × *D. nivalis* Norway (*D.n.* (N)), n = 13; and **(C)** *D.n.* (N) × *D.n.* (A), n = 21. Embryo stages: Unfertilized (Uf), Zygote (Zy), one-cell embryo proper (1C), two-cell embryo proper (2C),Octant (Oc), Dermatogen (De), Early globular (Eg), Mid globular (Mg) and Late globular (Lg). Endosperm stages: Unfertilized (Uf), Free floating (Ff), Syncytium (Sy), Micropylar cellularization (Mi) and Peripheral cellularization (Pe). Outliers are shown as dots. Scale bar = 50 µm.

To further investigate the endosperm cellularization time-point, we generated transgenic *D. nivalis* (Grimsdalen, Norway), a population that showed the same seed developmental progression as the *D. nivalis* (Juvasshytta, Norway) population used in the imprinting study (Supplementary Figure 6). Transgenic *proAT4G00220 >> H2A-GFP* (*TE1*) *D. nivalis* seeds were investigated from seven to 24 DAP (Figure 5, Supplementary Figure 7). In *A. thaliana,* the *TE1* reporter is expressed after initiation of endosperm cellularization and continues until the endosperm is consumed by the growing embryo (Bjerkan *et al.*, 2023; van Ekelenburg *et al.*, 2023). A highly similar expression pattern was observed in *D. nivalis*, showing weak *TE1-GFP* expression from 10 DAP, increasing in intensity at 12 DAP through 16 DAP before expression decrease again (Figure 5, Supplementary Figure 7). This demonstrates that *D. nivalis* seven DAP stages are well-suited for investigation of genomic imprinting before the onset of the developmental transition associated with endosperm cellularization.

**Figure 5:**
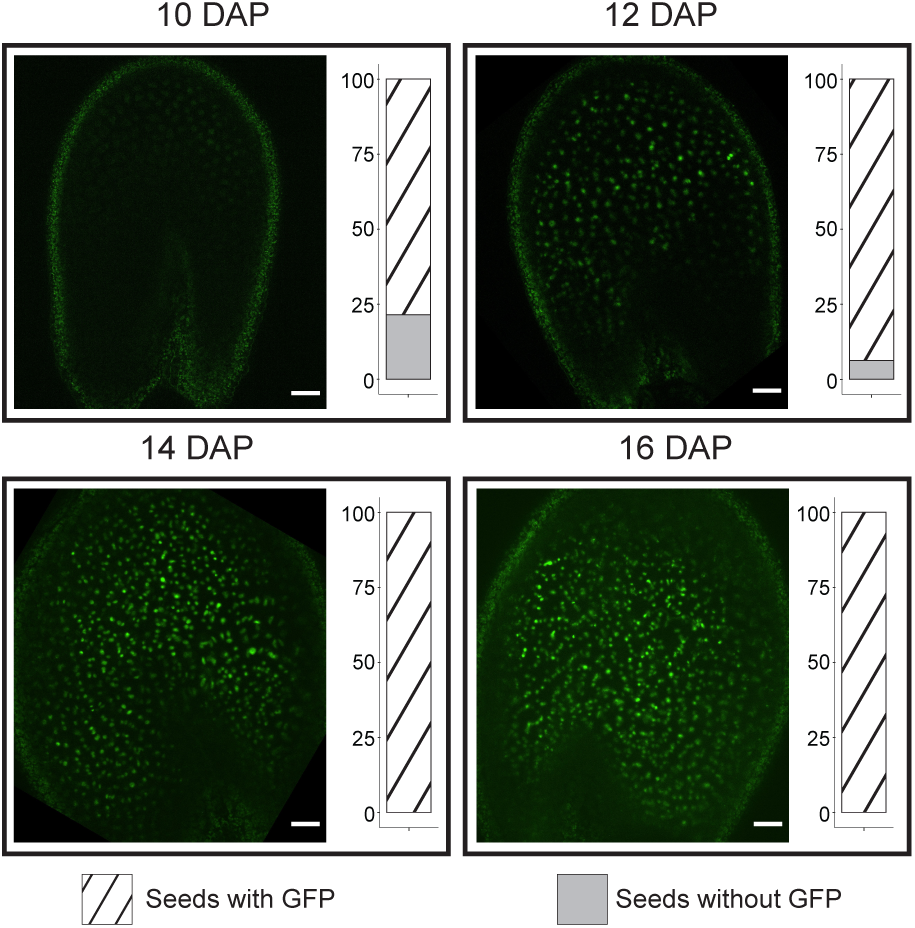
An endosperm cellularization reporter indicates endosperm cellularization in *Draba nivalis*. Confocal micrographs of *proAT4G00220>>H2A-GFP* (*TE1-GFP*) GFP expression in seeds of *D. nivalis* (population Grimsdalen, Norway) at different stages. In each panel the relative frequencies of seeds expressing *TE1-GFP* are indicated with diagonal stripes and seeds without expression are indicated as grey. Weak TE1-GFP expression is observed from 10 DAP. Biological replicates: 10 days after pollination (DAP) (n = 14), 12 DAP (n = 16), 14 DAP (n = 14) and 16 DAP (n = 18). Scale bar = 50 µm.

### Genomic imprinting analysis in *Draba nivalis* demonstrate large number of maternally expressed genes

We isolated RNA from the Alaska and Norway *D. nivalis* populations, in addition to their reciprocal hybrids from seeds at seven DAP. To establish good references for both parental populations, we mapped the parental reads to the *D. nivalis* Alaska transcriptome (Nowak *et al.*, 2020) and polished target sequences for each population with pilon (Walker *et al.*, 2014), which increased the mapping rate and quality of reference transcriptomes. To evaluate the differences between the Alaska and Norway *D. nivalis* populations, a differential expression analysis (DESeq2) (Love *et al.*, 2014) between the parents was performed on read counts generated by samstools (Danecek *et al.*, 2021). This analysis showed 3 265 differentially expressed genes in total, with 1830 genes upregulated in the Alaska × Norway cross and 1435 genes upregulated in the Norway × Alaska cross (Figure 6 A). Furthermore, a DESeq2 analysis on all samples, including the intraspecific hybrids, was performed in a similar manner to run a principal component analysis (PCA) (Blighe & Lun, 2023), which showed four distinct groups corresponding to the two parental lines and the two intraspecific hybrids (Figure 6 B). The parental lines were clearly separated by PC1, which explained 86.31% of the variation. However, PC2 explained only 6.14%, suggesting a substantial maternal bias.

**Figure 6:**
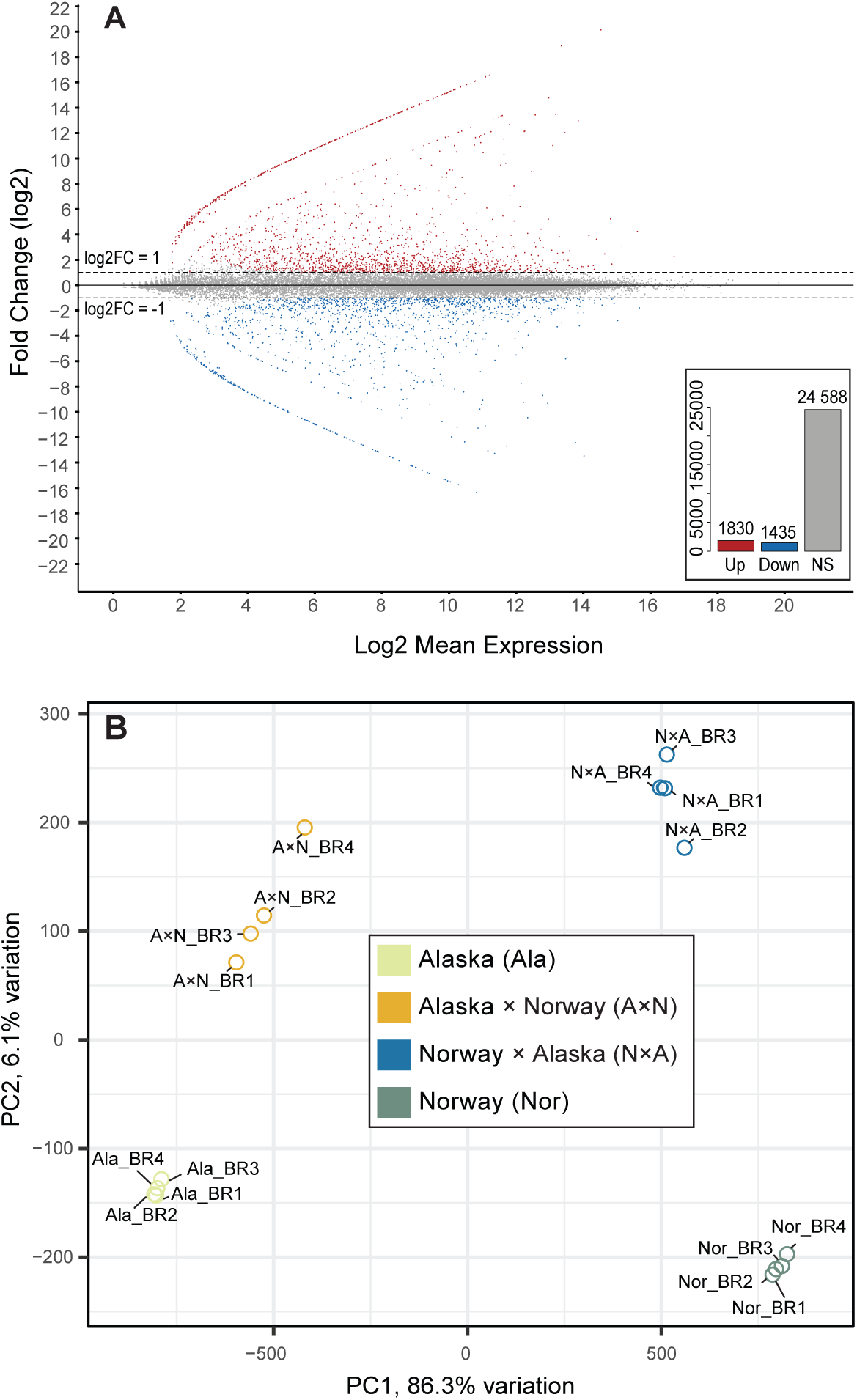
Differential gene expression between Alaska and Norway populations of *Draba nivalis* and homogeneity assessment of parental lines and intraspecific hybrids. **(A)** MA-plot showing the fold change differential expression between the Alaska and Norway populations. Differential expression is shown on the y-axis (log2). The x-axis shows the mean expression (log2). The analyses revealed 3265 genes to be significantly differentially expressed (inset box bottom right; red = upregulated, blue = downregulated, adjusted P-value < 0.05). Non-significantly regulated (NS) genes are depicted in gray. Dashed lines indicate log2 of 1 and -1. **(B)** Principal component analysis (PCA) based on gene expression of individual biological replicates from the two parental lines of *D. nivalis* from Alaska and Norway and their intraspecific hybrids. Crosses are indicated as maternal × paternal. Replicates from parental lines and intraspecific hybrids align homogeneously.

To identify imprinted genes, read pairs without sufficient single nucleotide polymorphisms (SNPs) or insertions/deletions (InDels), when mapping to the polished transcripts from both parents, were filtered out, retaining the informative reads (Hornslien *et al.*, 2019). Out of 33,557 genes, 11,319 had associated informative reads (Supplementary Data S5) and were selected for allele-specific differential expression analysis. Allele-specific total read counts for each biological replicate were counted and used to calculate statistical significance and fold change (FC) values (Supplementary Data S5). This dataset was used to identify genes with a parental bias and significant FC, which were considered imprinted (Supplementary Figure 8, Supplementary Data S5). The raw data indicated a large amount of MEGs (4468) and a small number of PEGs (16) and the high FC values for most of the dataset indicated a maternal bias (Supplementary Figure 8A).

Genes with very few read counts will achieve a sufficient FC value and significance, classifying it as an imprinted gene, even when differences are relatively small. This prompted the inclusion of additional filters before proceeding with the analyses. To address the problem with identification of imprinted genes with very few reads, genes with less than 40 reads mapped across all four replicates, excluding pseudocounts, were removed before the analysis. This decreased the number of PEGs extensively (10) (Supplementary Figure 8 B), resulting in a more reliable set of remaining PEGs (Table 1), whereas MEGs were only decreased slightly, from 4468 to 4411 (Supplementary Figure 8 B). To address the problem with an extensive maternal bias, we used a general seed coat (GSC) filter based on seed coat genes in *A. thaliana* (Hornslien *et al.*, 2019), which would be expected to contribute to maternal bias. Removing GSC genes from the dataset before the analysis resulted in a slight reduction of MEGs (4115), whereas the number of PEGs were not affected (Figure 7 A), but the distribution of FC values shows that the general maternal bias persisted (Figure 7 B). When considering the genomic ratios across the different seed tissues, a theoretical genomic ratio of 5:2 (maternal:paternal) is expected (embryo 1m:1p, seedcoat 2m:0p,endosperm 2m:1p) however, this estimation is an over-simplification, as it does not take cell number or gene regulation into account. In contrast, the *D. nivalis* dataset reveals a 9:1 (m:p) genomic ratio when considering only half of the maternal reads (Supplementary methods; Supplementary Data S5) and removing GSC genes, showing an exceedingly stronger maternal bias than what is expected from a theoretical genomic ratio.

**Figure 7:**
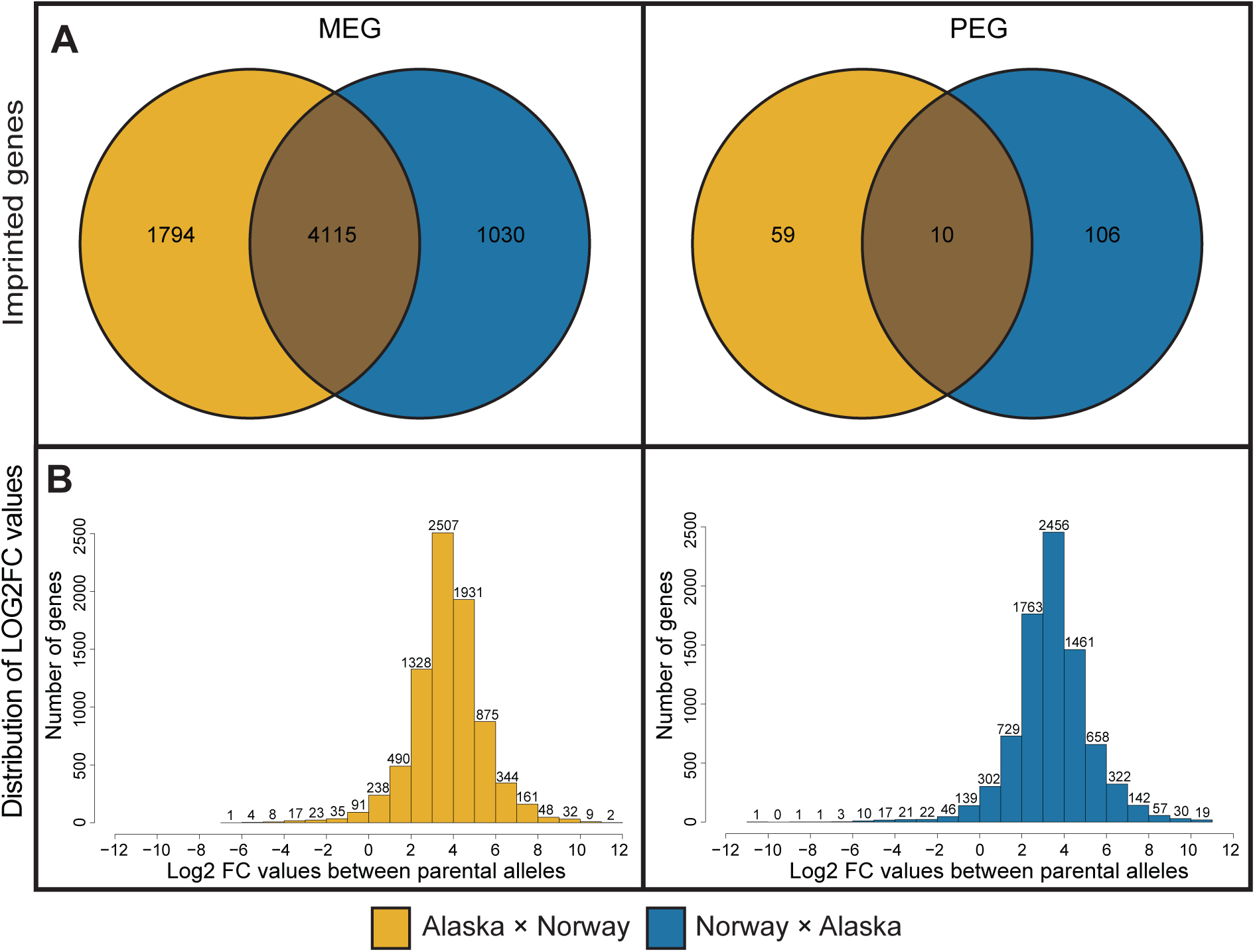
Distribution of maternally and paternally imprinted genes in *Draba nivalis*. **(A)** Venn diagrams showing MEGs and PEGs in reciprocal intraspecific *D. nivalis* crosses: Alaska × Norway (orange) and Norway × Alaska (blue). The union of reciprocal crosses are true imprinted genes, excluding genes that are overexpressed in a population specific manner. The representation is based on a filter allowing genes with more than 40 mapped reads (excluding pseudocounts), and excluding seed coat specific genes extrapolated from *A. thaliana* (Belmonte *et al.*, 2013). (B) Histograms showing the distribution of fold-change parental expression bias (log2, x-axis) in the reciprocal crosses Alaska × Norway (orange) and Norway × Alaska (blue). The absolute numbers of the fold-change classes are represented by the y-axis. Crosses are indicated as maternal × paternal.

**Table 1:** Paternally expressed genes (PEGs) identified in the intraspecific reciprocal cross between the Alaska and Norway populations of *Draba nivalis*. *Draba nivalis* PEGs identified in the genomic imprinting study, with the *Arabidopsis thaliana* ortholog (AGI), AGI associated gene description, log2 Fold Change (FC) values, FC values and adjusted P-value between maternal and paternal informative reads and the overlapping studies from Figure 9 B, which identified the same PEG.

| Gene | AGI | Gene description | Log2FC | FC<br>Pat/Mat<br>Ratio | Adjusted<br>P-value | Overlapping<br>study |
| --- | --- | --- | --- | --- | --- | --- |
| Dniva755 | AT5G02800 | Probable serine/threonine-protein kinase PBL7 | -5.3 | -39.7 | 1.07E-04 |  |
| Dniva6263 | AT3G49450 | F-box protein | -4.5 | -22.8 | 3.82E-05 | Del Toro-De León & Köhler, (2019) |
| Dniva4193 | AT2G43790 | Mitogen-activated protein kinase 6 | -3.0 | -8.1 | 5.48E-03 |  |
| Dniva32633 | AT2G24680 | B3 domain-containing protein REM12 | -4.2 | -18.6 | 6.55E-06 |  |
| Dniva22876 | AT1G34650 | Homeobox-leucine zipper protein HDG10 | -2.5 | -5.5 | 4.61E-04 | Wolff *et al.*, (2011); Hornslien *et al.*, (2019) |
| Dniva20625 | AT3G19600 | RNA polymerase II C-terminal domain phosphatase-like | -2.4 | -5.3 | 2.09E-03 |  |
| Dniva17206 | AT5G48670 | Agamous-like MADS-box protein AGL80 | -4.9 | -29.0 | 4.85E-05 |  |
| Dniva13170 | AT1G22220 | F-box protein AUF2 | -3.3 | -9.6 | 5.53E-04 |  |
| Dniva13061 | AT1G23320 | Tryptophan aminotransferase-related protein 1 | -3.5 | -11.2 | 1.54E-05 | Wolff *et al.*, (2011); Hornslien *et al.*, (2019) |
| Dniva12848 | AT5G50040 | Plant invertase/pectin methylesterase inhibitor superfamily protein | -3.5 | -11.6 | 4.10E-03 |  |

To better understand the pronounced maternal bias observed, we performed a gene ontology (GO) analysis of the identified MEGs. This resulted in enrichment of several biological processes involved in protein modification and folding, often including genes upregulated during stress involved in these processes (Figure 8). Interestingly, genes involved in these biological processes are highly duplicated and strongly expressed in *D. nivalis*, which is hypothesized to be an adaptation to the arctic environment (Nowak *et al.*, 2020).

**Figure 8:**
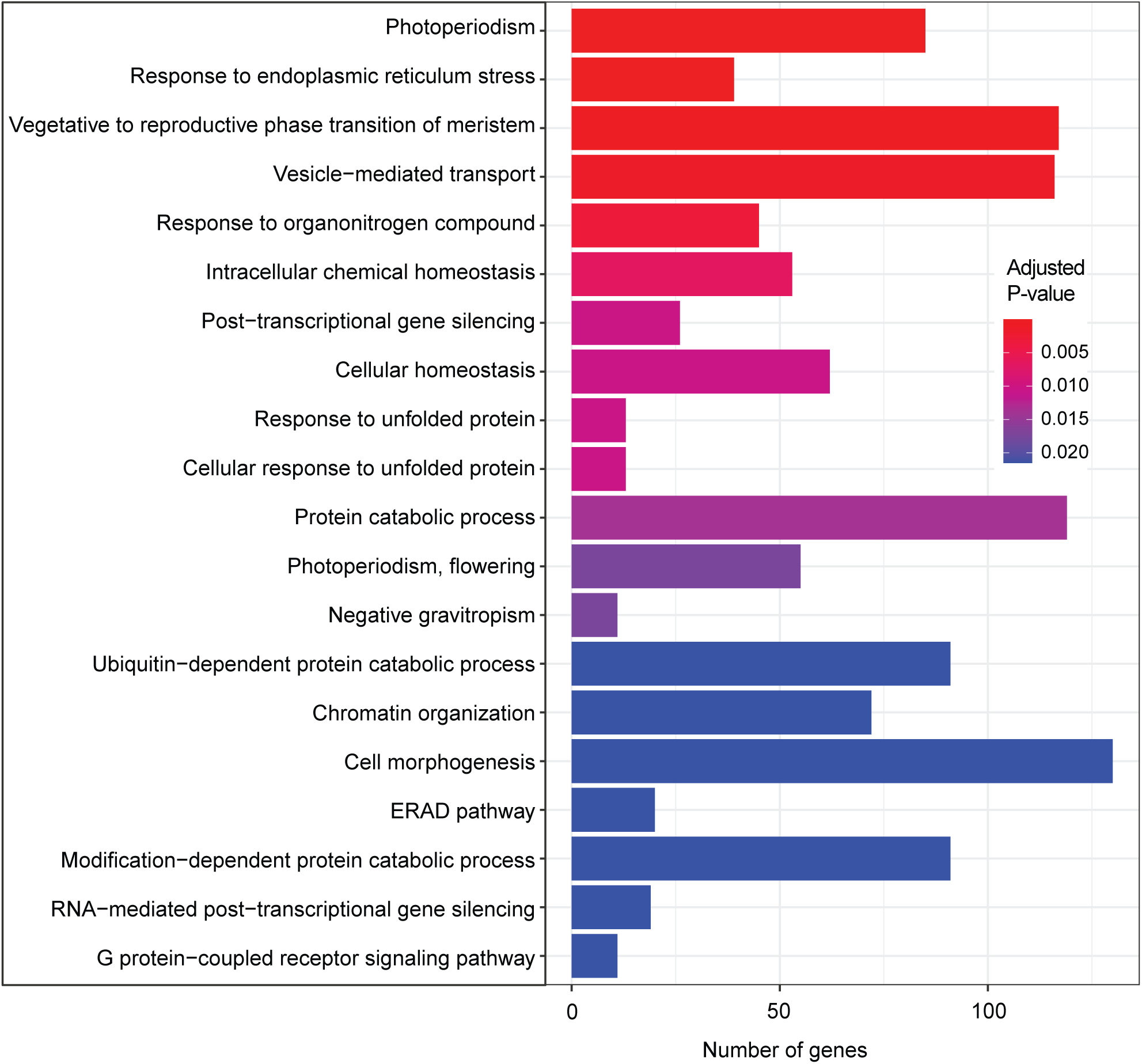
Gene ontology analysis of maternally expressed genes (MEGs) identified in *Draba nivalis.* MEGs from *D. nivalis* (cf. Figure 7A) that could be identified with a unique *Arabidopsis thaliana* ortholog were analyzed (Supplementary Data S4). Enriched biological GO processes are shown on the y-axis and the actual number of genes in each GO class is depicted by the x-axis. Adjusted p-values were calculated using the Benjamini-Hochberg method (Benjamini & Hochberg, 1995) and illustrated using a heatmap. Statistical significance (see inset box with heat map of p-values) is high when red and lower when blue

### Overlap of imprinting studies in Brassicaceae species

To assess the number of shared imprinted genes between *D. nivalis* and other Brassicaceae species, we conducted a comparative analysis. Previous studies on genomic imprinting in Brassicaceae have primarily focused on species within the supertribe Camelinodae, which potentially could have a higher overlap of imprinted genes than they have with the more distantly related species *D. nivalis*, belonging to the supertribe Arabodae. We found that out of the 4115 genes identified as MEGs in *D. nivalis*, 3429 had a clear ortholog in *A. thaliana*, facilitating comparison with other studies (Figure 9). Among these, 3107 showed no overlap with other studies, signifying that only 9.4% of all *D. nivalis* MEGs were also identified in one or more of the other Brassicaceae studies (Supplementary Table 3). The number of overlapping imprinted genes generally increased with larger datasets, showing the largest overlap with 151 MEGs in *A. thaliana* identified by (Del Toro-De León & Köhler, 2019)Figure 9 A). When comparing PEGs with other studies, we included all PEGs identified in either cross directions (Supplementary Data S5), excluding any population specific genes (Supplementary Data S5). Of the resulting 41 PEGs, 30 had no overlap with other studies (Figure 9 B, Supplementary Table 2). However, when examining the ten PEGs identified in both reciprocal intraspecific *D. nivalis* hybrids, three were discovered in one or more of the other Brassicaceae studies (Table 1), constituting an astounding 30% (Supplementary Table 3).

**Figure 9:**
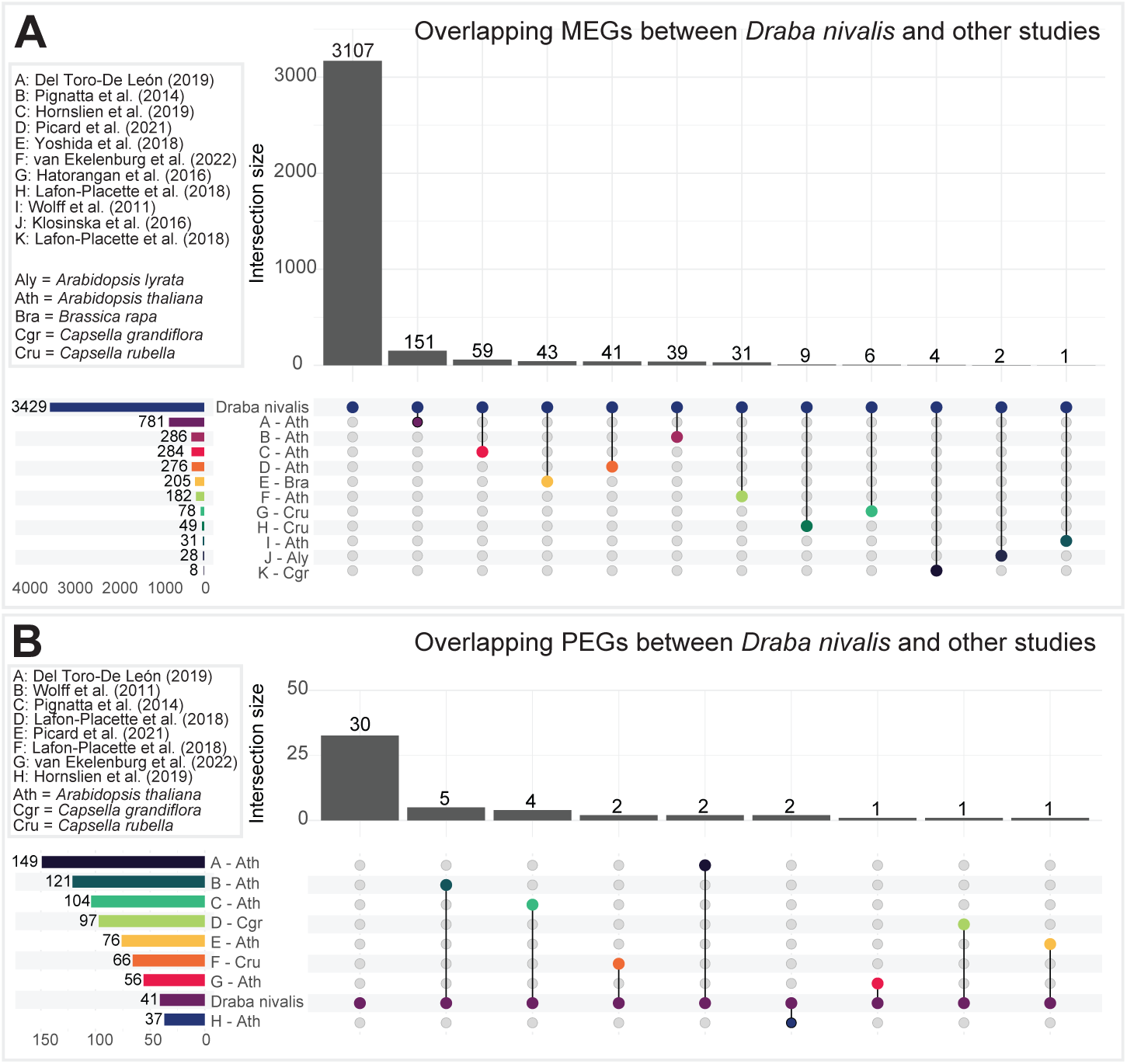
MEGs and PEGs in *Draba nivalis* previously identified in Brassicaceae species. The upper left box shows previous studies used in the comparison and species abbreviations. The lower left box shows the total number of MEGs/PEGs from each study. The intersection size histogramme indicates the number of genes overlapping with each study, with the first intersection bar being the number of *D. nivalis* genes with no overlap. **(A)** MEGs overlapping between *D. nivalis* and species from 11 other studies (A-K), including *Arabidopsis thaliana, A. lyrata, Brassica rapa, Capsella grandiflora* and *C. rubella*. **(B)** PEGs overlapping between *D. nivalis* and species from eight other studies (A-H), including *A. thaliana, C. grandiflora* and *C. rubella*. The PEGs included in the comparison considers all identified *D. nivalis* PEGs identified regardless of cross direction (Fig. 7), after filtering away genes that are biased either in the Alaska × Norway cross or the Norway × Alaska cross (Fig. 6A).

We performed a new GO-analysis using only the MEGs overlapping between *D. nivalis* and other Brassicaceae species (Figure 9 A, Figure 10) and compared with our previous GO-analysis using all MEGs from *D. nivalis* (Figure 8). Interestingly, the most significant group of MEGs is involved in syncytium formation, which is an important stage during early endosperm development. This group contained four expansin genes, AtEXPA1, AtEXPA6, AtEXPA10, and AtEXPA15, in addition to Inositol oxygenase 1, which are all involved in abiotically triggered syncytium formation in the *A. thaliana* root (Wieczorek *et al.*, 2006; Siddique *et al.*, 2009). However, no study so far has researched their involvement in endosperm syncytium, although expression of all genes is found within the seed (Mergner *et al.*, 2020). Furthermore, there were GO-groups involving cell wall biogenesis, modification and organization, which could be involved in the cellularization process of the endosperm. There was a strong difference in the identified GO-terms using all MEGs from *D. nivalis* and using only MEGs overlapping with species from other studies (Figure 8, Figure 9 A), emphasizing a general difference between *D. nivalis* and other Brassicaceae species.

**Figure 10:**
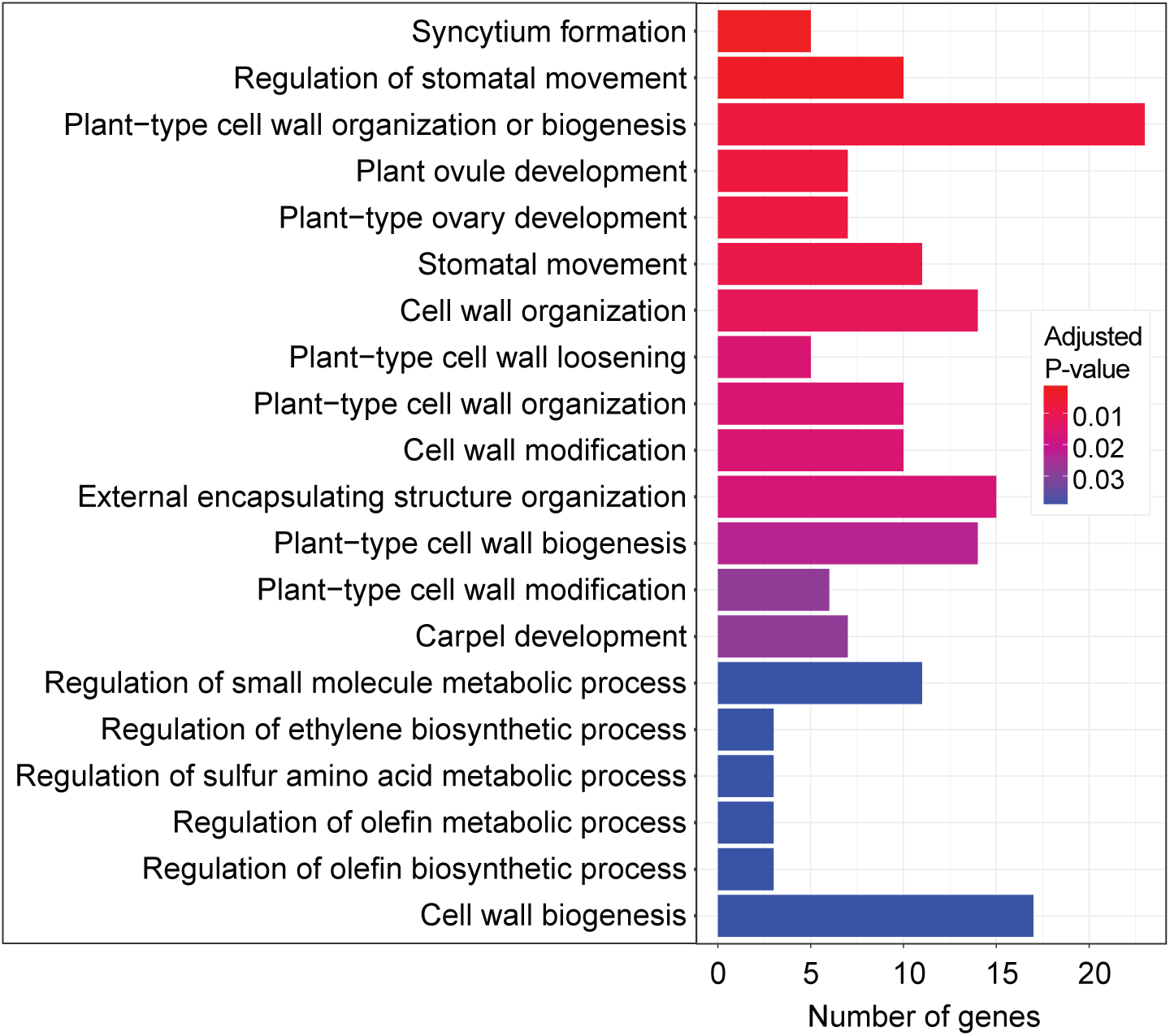
Gene ontology (GO) analysis of MEGs identified in *Draba nivalis* and other Brassicaceae species. The analysis contains enriched biological processes obtained from overlapping MEGs between *D. nivalis* and Brassicaceae species from other studies (Figure 9 A). MEGs from *D. nivalis* (cf. Figure 7A) that could be identified with a unique *Arabidopsis thaliana* ortholog were analyzed (Supplementary Data S4). Enriched biological GO processes are shown on the y-axis and the actual number of genes in each GO class is depicted by the x-axis. Adjusted p-values were calculated using the Benjamini-Hochberg method (Benjamini & Hochberg, 1995) and illustrated using a heatmap. Statistical significance (see inset box with heat map of p-values) is high when red and lower when blue. Several genes involved in syncytium formation and cell wall regulation are regulated by genomic imprinting.

## Discussion

Although there is previous report that pre- and/or post-zygotic barriers might have affected seed set and seed viability in crosses between the three studied species of *Draba* (Mulligan, 1974; Brochmann *et al.*, 1993), we found no such evidence in our study, based on extensive crossing experiments between populations from different geographic areas of these three highly selfing diploid species. Rather, our results provide strong support for the hypothesis that endosperm-based hybrid barriers do not evolve in highly selfing species because of absence or low levels of parental conflict. Several factors have been shown to increase or decrease the strength of hybrid barriers by affecting endosperm cellularization time point during hybrid seed development. This includes both genetic and abiotic factors, such as genomic imprinting, temperature, single-gene mutations, ploidy and cross direction of the parental individuals (Lafon-Placette *et al.*, 2017; Bjerkan *et al.*, 2020, 2023). However, mating system (selfing versus outbreeding) may also strongly influence the development of endosperm-based hybrid barriers (Bjerkan *et al.*, 2020, 2023; Petrén *et al.*, 2023) as well as other types of barriers such as hybrid sterility (Marie-Orleach *et al.*, 2022, 2023). Whereas both inter- and intraspecific F_1_ hybrids involving these three *Draba* species frequently are fully or partly sterile (Mulligan, 1974; Grundt *et al.*, 2006), with accumulation of hybrid incompatibilities most likely accelerated by their high selfing rates (Marie-Orleach *et al.*, 2023), our study suggests that selfing has the opposite effect on development of endosperm-based hybrid barriers by reducing parental conflict.

Mating system appears to influence not only hybrid barriers, but also genomic imprinting. Outbreeding species, in which the levels of parental conflict are high, show an increase in number of PEGs compared to selfing species, while not affecting the number of MEGs (Klosinska *et al.*, 2016; Lafon-Placette *et al.*, 2018). Interestingly, our investigation of genomic imprinting in *D. nivalis* identified numerous MEGs and very few PEGs. Despite the few PEGs identified, several of them have been found in other Brassicaceae species, which is not the case for the MEGs identified in this study. A generally low level of conservation of genomic imprinting, even between closely related species, is a well-known phenomenon but not entirely understood. The selfing strategy of *D. nivalis* cannot alone explain the genomic imprinting trends we observed, rather, the enriched GO-terms of the MEGs suggest that the harsh arctic environment affects genomic imprinting in this species. Based on the results from this study, we propose that in addition to the mating system, environmental adaptations may exert significant effects on genomic imprinting dynamics.

### Absence of endosperm hybrid barriers may be explained by low parental conflict in highly selfing species

We found no endosperm-based hybridization barrier in interspecific crosses between *D. fladnizensis*, *D. nivalis* and *D. subcapitata*. Even though this contradicts previous reports (Mulligan, 1974; Brochmann *et al.*, 1993), it is not unexpected when considering they are highly selfing species. Outbreeding species will exhibit strong parental conflict, because the maternal individuals may be fertilized by other paternal individuals. For the maternal parent it will be more beneficial with an equal resource allocation to the developing embryos, whereas each paternal counterpart will benefit from increased resource allocation to its own progeny alone. However, in selfing plants, selection for parental conflict in resource allocation is not necessary. This is the basis for the weak inbreeder/strong outbreeder (WISO) hypothesis, which predicts that differences in parental conflict between outbreeding and selfing species will cause an imbalance in resource allocation to the embryo (Brandvain & Haig, 2005, 2018; Raunsgard *et al.*, 2018). In Brassicaceae species, which have nuclear endosperm development, such imbalance may cause a precocious or delayed cellularization of the endosperm, which further affects embryo viability (Lafon-Placette & Köhler, 2014; Lafon-Placette *et al.*, 2017, 2018; Bjerkan *et al.*, 2023). Since all the examined *Draba* species are highly selfing (Brochmann, 1993), one would expect low parental conflict and consequently no endosperm hybrid barrier in accordance with our results.

### Low parental conflict alone is insufficient to explain maternal bias during seed development

In our study of genomic imprinting in *D. nivalis*, we found an expected decrease in PEGs, accompanied by an unexpected and substantial increase in MEGs, suggesting a strong maternal bias. The number and expression level of PEGs have been found to be higher in outbreeding species than in selfing species. In *Arabidopsis*, more PEGs were identified in the outbreeder *A. lyrata* than in the selfing *A. thaliana* (Klosinska *et al.*, 2016), and in *Capsella*, more PEGs were identified in the outbreeder *C. grandiflora* than in the recent selfer *C. rubella* and the ancient selfer *C. orientalis* (Lafon-Placette *et al.*, 2018). In contrast to our findings, the number and expression of MEGs did not differ in these studies, showing that the mating system has an effect on genomic imprinting. Although a maternal bias is considered beneficial in selfing species as it allows for equal resource allocation across all progenies (Haig & Westoby, 1991; Brandvain & Haig, 2005), such an extreme maternal bias as we found in *D. nivalis* has not been reported for other selfing species, suggesting that other factors may play a role in addition to the selfing strategy. This is supported by the results from the GO-term analysis, where the enriched GO-terms based on all MEGs or only MEGs overlapping with other studies, differed completely. This suggests that the high abundance of MEGs may result from specific adaptations in *D. nivalis*. Interestingly, the GO-terms enriched for all MEGs were also reported as enriched in *D. nivalis* by Nowak et al. (2020). They found that these include genes that are highly duplicated and strongly expressed, hypothesizing that this could be an adaptation to the extreme arctic environment (Nowak *et al.*, 2020). Accordingly, the selfing strategy, together with adaptations to the arctic environment, may be responsible for the observed strong maternal bias in *D. nivalis*. Maternal bias could be beneficial in harsh environments, as it allows for adaptation to the maternal environment, which has been reported to have great importance for seed germination (Veselá *et al.*, 2021).

### Reduced number of paternally expressed genes overlap with other Brassicaceae species

There are only a few reports of overlap in imprinted genes across species, and several explanations have been proposed. One explanation states that using different species or ecotypes will necessarily result in different genes being analyzed, because the analysis relies on identification of SNPs or InDels, which vary across the parental individuals (Zhang *et al.*, 2011; Pignatta *et al.*, 2014; van Ekelenburg *et al.*, 2023). In spite of the large number of MEGs identified in this study, we found few overlaps with other studies. Only 9.4% of the MEGs identified in *D. nivalis* overlapped with other Brassicaceae studies (Supplementary Table 3). However, some of the overlapping genes were enriched for GO-terms involving syncytium development and cell wall modification, showing that there is some conservation in genomic imprinting across different species. In contrast, among the very few PEGs identified, as many as 30% were overlapping with other studies, suggesting that although the number of PEGs are severely reduced, the most important genes keep their imprinting status. Compared to the other Brassicaceae species with identified imprinted genes, *D. nivalis* is, together with *B. rapa*, the most distantly related; all other species belong to supertribe Camelinodae. This makes it both harder to find a good *A. thaliana* ortholog and it increases the likelihood for differences in genomic imprinting. More overlap in both MEGs and PEGs are found between the Camelinodae species than between *D. nivalis* and *B. rapa* (Supplementary Table 3). However, for *B. rapa* we find a higher overlap in MEGs to the other Brassicaceae species (33.7%) compared to overlaps in PEGs (18.2%), which is the inverse trend of what we find for *D. nivalis*. Furthermore, a study in *A. thaliana* using endosperm nuclei sorting (van Ekelenburg *et al.*, 2023), resulted in approximately 33.9% overlap in PEGs to the other studies, which is very close to what is found in *D. nivalis*. Altogether this suggests that the imprinting status of the few remaining PEGs in *D. nivalis* is of importance for seed development. It should be noted that the overlap in PEGs between the selfing *C. rubella* and the other studies is twice as high as what is found for the outbreeding *C. grandiflora* (Hatorangan *et al.*, 2016; Lafon-Placette *et al.*, 2018), showing that the PEGs preserved in *C. rubella* are more likely to have an important and conserved role in seed development. By using information on mating systems and potential environmental adaptations, we can more precisely pinpoint which imprinted genes have conserved importance across more distantly related species. Furthermore, by comparing species with similar mating strategies and environmental adaptations, a higher overlap in imprinted genes may be found.

In summary, we have demonstrated that an endosperm-based post-zygotic hybridization barriers are absent in interspecies hybrid *Draba* seeds, supporting low parental conflict. In a *D. nivalis* genomic imprinting study we report a high number of maternally expressed genes (MEGs) and low numbers of paternally expressed genes (PEGs), suggesting rapid evolution of MEGs and loss of PEGs in a mating system with low parental conflict, and proposing that selfing arctic species may exhibit a generally stronger maternal expression bias as an adaptive mechanism to efficiently cope with the arctic environment.

## Acknowledgements

We thank Jason R. Miller for generous bioinformatics advice and the staff at the UiO:PlantLab Marit Langrekken, Ingrid Johansen and Vegard Iversen for maintenance of plants.

## Supplementary Tables

**Table S1:** Coordinates, origin and references of species and population used in this study. Collector abbreviations: Reidar Elven (RE), Hanne Hegre Grundt (HHG), Anne Krag Brysting (AKB), Christian Brochmann (CB), Jorun Nyhlen (JN), Pernille Bronken Eidesen (PBE) and Siri Kjølner (SK).

| Species | Population | ID | Coordinates | Origin | Collection year | Collector | Reference |
| --- | --- | --- | --- | --- | --- | --- | --- |
| *Draba nivalis* | Alaska | 8-4 | N 63.045,<br>E -147.201 | USA, Alaska,<br>Alaska range,<br>Clearwater Mts,<br>Waterfall Creek<br>W | 1998 | RE, HHG | Grundt *et al.*, (2006) |
| *Draba nivalis* | Greenland | AKB17-3 | N 69.730555,<br>E -51.933606 | Greenland,<br>Disko,<br>Mudderbugten | 2017 | AKB | This manuscript |
| *Draba nivalis* | Norway | AKB17-2 | N 61.674307,<br>E 8.368491 | Norway,<br>Oppland, Lom,<br>Juvasshytta | 2017 | AKB | This manuscript |
| *Draba nivalis* | Norway | 45-5 | N 62.06159,<br>E 9.5403 | Grimsdalen,<br>Rondane | 1999 | HHG | Grundt *et al.*, (2006) |
| *Draba fladnizensis* | Alaska | 4-1 | N 65.5,<br>E -145.354 | USA, Alaska,<br>Eagle summit S | 1998 | HHG, RE | Grundt *et al.*, (2006) |
| *Draba fladnizensis* | Greenland | 74-12 | N 70.733,<br>E -22.667 | Greenland,<br>East Greenland,<br>Jameson land,<br>Constable Point | 1999 | CB, JN,<br>PBE, SK | Grundt *et al.*, (2006) |
| *Draba fladnizensis* | Norway | AKB17-2 | N 61.674307,<br>E 8.368491 | Norway,<br>Oppland, Lom,<br>Juvasshytta | 2017 | AKB | This manuscript |
| *Draba subcapitata* | Greenland | 81-12 | N 70.7135, E<br>-22.695889 | Greenland,<br>East Greenland,<br>Jameson land,<br>Harris Fjeld | 1999 | CB, JN,<br>PBE, SK | Grundt *et al.*, (2006) |

**Table S2:** Paternally expressed genes (PEGs) identified in either reciprocal cross between the Alaska and Norway populations of *Draba nivalis*. *Draba nivalis* PEGs identified in the genomic imprinting study of *D. nivalis*, with the *Arabidopsis thaliana* ortholog (AGI), AGI associated gene description, log2 Fold Change (FC) values, FC values given as paternal/maternal ratio and Adjusted P-value between maternal and paternal informative reads, and the overlapping studies identifying the same PEG.

| Gene | AGI | Gene description | Log2FC | FC<br>Paternal/<br>Maternal<br>Ratio | Adjusted P-value | Overlapping study |
| --- | --- | --- | --- | --- | --- | --- |
| Dniva12293 | AT1G28630 | Transcriptional regulator EFH1-like protein | -2.0 | 4.1 | 1.71E-03 | Pignatta *et al.*, (2014); Lafon-Placette *et al.*, (2018); van Ekelenburg *et al.*, (2023) |
| Dniva15348 | AT1G07980 | DNA polymerase II subunit B3-1 | -3.4 | 10.3 | 2.26E-05 |  |
| Dniva15369 | AT2G42470 | MATH domain and coiled-coil domain-containing protein | -2.3 | 4.8 | 3.65E-04 |  |
| Dniva16002 | AT1G02790 | Exopolygalacturonase clone GBGE184 | -2.6 | 6.0 | 3.86E-02 | Wolff *et al.*, (2011) |
| Dniva195 | AT1G11900 | Pentatricopeptide repeat-containing protein | -2.7 | 6.5 | 6.16E-05 |  |
| Dniva31779 | AT5G45880 | Ole e I (Main olive allergen)-like protein | -2.8 | 6.9 | 2.05E-02 |  |
| Dniva32251 | AT4G11720 | Protein HAPLESS 2 | -1.5 | 2.8 | 1.43E-02 |  |
| Dniva32999 | NA |  | -3.5 | 11.0 | 1.21E-05 |  |
| Dniva6691 | NA |  | -2.3 | 4.8 | 2.39E-03 |  |
| Dniva7010 | AT2G34555 | Gibberellin 2-beta-dioxygenase 3 | -1.3 | 2.5 | 2.53E-02 |  |
| Dniva7227 | AT2G35765 | Uncharacterized protein | -2.2 | 4.6 | 1.59E-02 | Wolff *et al.*, (2011) |
| Dniva8746 | AT1G77020 | DNAJ heat shock N-terminal domain-containing protein | -1.8 | 3.4 | 3.54E-02 |  |
| Dniva10631 | AT1G57660 | Large ribosomal subunit protein eL21x/eL21w | -5.6 | 46.9 | 3.03E-08 |  |
| Dniva11491 | AT1G61320 | FBD-associated F-box protein | -3.4 | 10.9 | 3.60E-04 | Pignatta *et al.*, (2014) |
| Dniva12159 | AT1G29760 | Seipin-2 | -1.9 | 3.7 | 3.92E-03 |  |
| Dniva12270 | AT1G29080 | Papain family cysteine protease | -2.2 | 4.7 | 3.45E-03 |  |
| Dniva1334 | AT2G47430 | Histidine kinase CKI1 | -3.6 | 12.5 | 6.67E-05 |  |
| Dniva16372 | AT5G23740 | Small ribosomal subunit protein uS17x | -1.4 | 2.7 | 2.74E-03 |  |
| Dniva18999 | AT3G18910 | F-box protein ETP2 | -1.8 | 3.5 | 8.49E-03 | Lafon-Placette *et al.*, (2018) |
| Dniva20432 | AT5G50070 | Plant invertase/pectin methylesterase inhibitor superfamily protein | -2.1 | 4.2 | 2.01E-03 |  |
| Dniva22683 | NA |  | -1.1 | 2.1 | 1.81E-03 |  |
| Dniva24907 | AT5G60760 | P-loop NTPase domain-containing protein LPA1 homolog 1 | -1.8 | 3.5 | 3.54E-03 | Pignatta *et al.*, (2014) |
| Dniva26666 | AT4G39000 | Endoglucanase 23 | -1.2 | 2.3 | 1.99E-03 |  |
| Dniva2839 | AT3G46840 | Subtilisin-like protease SBT4.5 | -1.7 | 3.3 | 1.38E-04 |  |
| Dniva28976 | AT4G23120 | Bromo-adjacent homology (BAH) domain-containing protein | -3.1 | 8.3 | 1.14E-05 | Pignatta *et al.*, (2014) |
| Dniva29121 | AT4G23800 | High mobility group B protein 6 | -3.7 | 12.6 | 1.15E-07 |  |
| Dniva29651 | AT1G11593 | Plant invertase/pectin methylesterase inhibitor superfamily protein | -1.6 | 3.0 | 1.24E-04 |  |
| Dniva30935 | AT2G32275 | Expressed protein | -2.6 | 6.2 | 7.35E-05 | Wolff *et al.*, (2011); Del Toro-De León & Köhler, (2019); Picard *et al.*, (2021) |
| Dniva33135 | AT5G39850 | Small ribosomal subunit protein uS4y | -1.7 | 3.2 | 2.37E-03 |  |
| Dniva4832 | AT4G05320 | Polyubiquitin 10 | -1.4 | 2.7 | 4.66E-02 |  |
| Dniva5365 | AT1G28500 | Putative UPF0725 protein | -1.6 | 3.1 | 6.16E-03 |  |
| Dniva5913 | AT4G13780 | Methionine--tRNA ligase, cytoplasmic | -5.9 | 59.7 | 1.43E-08 |  |
| Dniva6270 | AT5G06500 | AGAMOUS-like 96 | -2.8 | 7.0 | 6.43E-05 |  |
| Dniva7113 | AT2G34990 | RING-H2 finger protein ATL38 | -1.9 | 3.8 | 1.07E-02 |  |

**Table S3:**
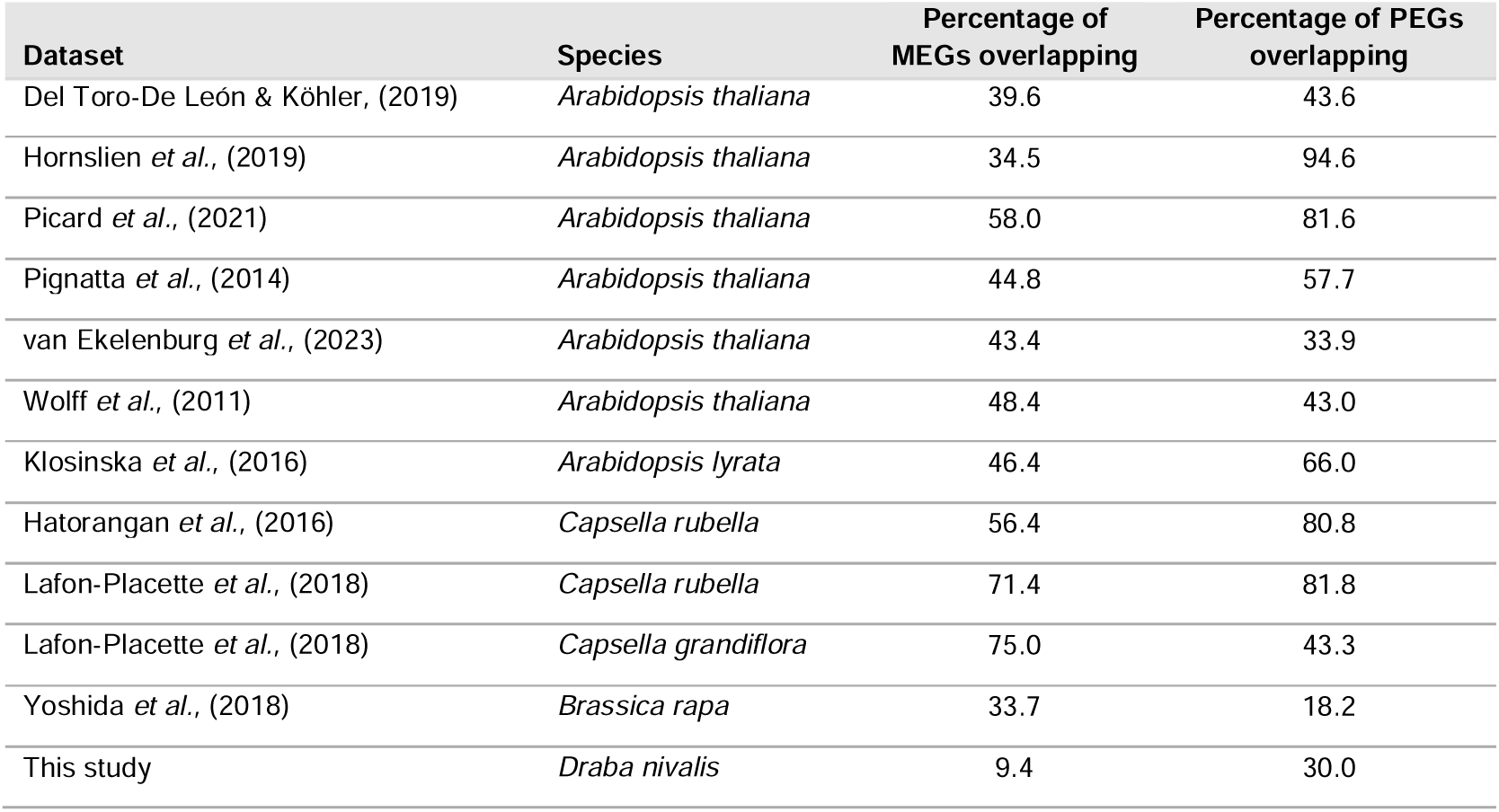
Overlap between imprinted genes in Brassicaceae species. The table shows imprinting studies in Brassicaceae species and the percentage of maternally expressed genes (MEGs) and paternally expressed genes (PEGs) which overlap with MEGs and PEGS found in other studies.

## References

Al-Shehbaz IA, Beilstein MA, Kellogg EA. 2006. Systematics and phylogeny of the Brassicaceae (Cruciferae): an overview. Plant systematics and evolution = Entwicklungsgeschichte und Systematik der Pflanzen 259: 89–120.

AU-Lindsey BE III, AU-Rivero L, AU-Calhoun CS, AU-Grotewold E, AU-Brkljacic J. 2017. Standardized Method for High-throughput Sterilization of Arabidopsis Seeds. Journal of visualized experiments: JoVE: e56587.

Bailey CD, Koch MA, Mayer M, Mummenhoff K, O’Kane SL Jr, Warwick SI, Windham MD, Al-Shehbaz IA. 2006. Toward a Global Phylogeny of the Brassicaceae. Molecular biology and evolution 23: 2142–2160.

Barrett SCH. 2002. The evolution of plant sexual diversity. Nature reviews. Genetics 3: 274–284.

Batista RA, Köhler C. 2020. Genomic imprinting in plants—revisiting existing models. Genes & development 34: 24–36.

Belmonte MF, Kirkbride RC, Stone SL, Pelletier JM, Bui AQ, Yeung EC, Hashimoto M, Fei J, Harada CM, Munoz MD, et al. 2013. Comprehensive developmental profiles of gene activity in regions and subregions of the Arabidopsis seed. Proceedings of the National Academy of Sciences of the United States of America 110: E435–44.

Benjamini Y, Hochberg Y. 1995. Controlling the false discovery rate: a practical and powerful approach to multiple testing. Journal of the Royal Statistical Society. Series B, Statistical methodology 57: 289–300.

Birchler JA. 1993. DOSAGE ANALYSIS OF MAIZE ENDOSPERM DEVELOPMENT. Annual review of genetics 27: 181–204.

Bjerkan KN, Alling RM, Myking IV, Brysting AK, Grini PE. 2023. Genetic and environmental manipulation of Arabidopsis hybridization barriers uncovers antagonistic functions in endosperm cellularization. Frontiers in plant science 14.

Bjerkan KN, Hornslien KS, Johannessen IM, Krabberød AK, van Ekelenburg YS, Kalantarian M, Shirzadi R, Comai L, Brysting AK, Bramsiepe J, et al. 2020. Genetic variation and temperature affects hybrid barriers during interspecific hybridization. The Plant journal: for cell and molecular biology 101: 122–140.

Blighe K, Lewis M, Lun A, Blighe MK. 2019. Package ‘PCAtools.’ *REAGENT or RESOURCE SOURCE IDENTIFIER Deposited data GitHub repository with code and data for analysis GitHub* https://github.com/CharlotteEPage/NFI_Disease_Ecology Raw data tables Mendeley https://data.mendeley.com/datasets/y869bhhmzr/1 Software and algorithms R version 4: 9–11.

Blighe K, Lun A. 2023. PCAtools: PCAtools: Everything Principal Components Analysis.

Brandvain Y, Haig D. 2005. Divergent Mating Systems and Parental Conflict as a Barrier to Hybridization in Flowering Plants. The American naturalist 166: 330–338.

Brandvain Y, Haig D. 2018. Outbreeders pull harder in a parental tug-of-war. Proceedings of the National Academy of Sciences of the United States of America 115: 11354–11356.

Brochmann C. 1993. Reproductive strategies of diploid and polyploid populations of arctic Draba (Brassicaceae). Plant systematics and evolution = Entwicklungsgeschichte und Systematik der Pflanzen 185: 55–83.

Brochmann C, Borgen L, Stedje B. 1993. Crossing relationships and chromosome numbers of Nordic populations of Draba (Brassicaceae), with emphasis on the D. alpina complex. Nordic journal of botany 13: 121–147.

Bushell C, Spielman M, Scott RJ. 2003. The basis of natural and artificial postzygotic hybridization barriers in Arabidopsis species. The Plant cell 15: 1430–1442.

Chen H, Boutros PC. 2011. VennDiagram: a package for the generation of highly-customizable Venn and Euler diagrams in R. BMC bioinformatics 12: 35.

Clough SJ, Bent AF. 1998. Floral dip: a simplified method for Agrobacterium-mediated transformation of Arabidopsis thaliana. The Plant journal: for cell and molecular biology 16: 735–743.

Cooper DC, Brink RA. 1942. The Endosperm as a Barrier to Interspecific Hybridization in Flowering Plants. Science 95: 75–76.

Cooper DC, Brink RA. 1945. SEED COLLAPSE FOLLOWING MATINGS BETWEEN DIPLOID AND TETRAPLOID RACES OF LYCOPERSICON PIMPINELLIFOLIUM. Genetics 30: 376–401.

Cornejo P, Camadro EL, Masuelli RW. 2012. Molecular bases of the postzygotic barriers in interspecific crosses between the wild potato species Solanum acaule and Solanum commersonii. Genome / National Research Council Canada = Genome / Conseil national de recherches Canada 55: 605–614.

Danecek P, Bonfield JK, Liddle J, Marshall J, Ohan V, Pollard MO, Whitwham A, Keane T, McCarthy SA, Davies RM, et al. 2021. Twelve years of SAMtools and BCFtools. GigaScience 10.

Del Toro-De León G, Köhler C. 2019. Endosperm-specific transcriptome analysis by applying the INTACT system. Plant reproduction 32: 55–61.

Dong X, Luo H, Bi W, Chen H, Yu S, Zhang X, Dai Y, Cheng X, Xing Y, Fan X, et al. 2023. Transcriptome-wide identification and characterization of genes exhibit allele-specific imprinting in maize embryo and endosperm. BMC plant biology 23: 470.

Dziasek K, Simon L, Lafon-Placette C, Laenen B, Wärdig C, Santos-González J, Slotte T, Köhler C. 2021. Hybrid seed incompatibility in Capsella is connected to chromatin condensation defects in the endosperm. PLoS genetics 17: e1009370.

van Ekelenburg YS, Hornslien KS, Van Hautegem T, Fendrych M, Van Isterdael G, Bjerkan KN, Miller JR, Nowack MK, Grini PE. 2023. Spatial and temporal regulation of parent-of-origin allelic expression in the endosperm. Plant physiology 191: 986–1001.

Elven R, Arnesen G, Alsos IG, Sandbakk BE. 2020. Svalbardflora second edition. https://svalbardflora.no/.

Emms DM, Kelly S. 2019. OrthoFinder: phylogenetic orthology inference for comparative genomics. Genome biology 20: 238.

Flores-Vergara MA, Oneal E, Costa M, Villarino G, Roberts C, De Luis Balaguer MA, Coimbra S, Willis J, Franks RG. 2020. Developmental analysis of Mimulus seed transcriptomes reveals functional gene expression clusters and four imprinted, endosperm-expressed genes. Frontiers in plant science 11: 132.

Florez-Rueda AM, Paris M, Schmidt A, Widmer A, Grossniklaus U, Städler T. 2016. Genomic imprinting in the endosperm is systematically perturbed in abortive hybrid tomato seeds. Molecular biology and evolution 33: 2935–2946.

Grini PE, Jürgens G, Hülskamp M. 2002. Embryo and endosperm development is disrupted in the female gametophytic capulet mutants of Arabidopsis. Genetics 162: 1911– 1925.

Grundt HH, Kjølner S, Borgen L, Rieseberg LH, Brochmann C. 2006. High biological species diversity in the arctic flora. Proceedings of the National Academy of Sciences 103: 972–975.

Grundt HH, Popp M, Brochmann C, Oxelman B. 2004. Polyploid origins in a circumpolar complex in Draba (Brassicaceae) inferred from cloned nuclear DNA sequences and fingerprints. Molecular phylogenetics and evolution 32: 695–710.

Gustafsson ALS, Gussarova G, Borgen L, Ikeda H, Antonelli A, Marie-Orleach L, Rieseberg LH, Brochmann C. 2022. Rapid evolution of post-zygotic reproductive isolation is widespread in Arctic plant lineages. Annals of botany 129: 171–184.

Gustafsson ALS, Skrede I, Rowe HC, Gussarova G, Borgen L, Rieseberg LH, Brochmann C, Parisod C. 2014. Genetics of Cryptic Speciation within an Arctic Mustard, Draba nivalis. PloS one 9: e93834.

Haig D, Westoby M. 1991. Genomic Imprinting in Endosperm: Its Effect on Seed Development in Crosses between Species, and between Different Ploidies of the Same Species, and Its Implications for the Evolution of Apomixis. *Philosophical transactions of the Royal Society of London. Series B*, Biological sciences 333: 1–13.

Hatorangan MR, Laenen B, Steige KA, Slotte T, Köhler C. 2016. Rapid Evolution of Genomic Imprinting in Two Species of the Brassicaceae. The Plant cell 28: 1815–1827.

Hendriks KP, Kiefer C, Al-Shehbaz IA, Bailey CD, Hooft van Huysduynen A, Nikolov LA, Nauheimer L, Zuntini AR, German DA, Franzke A, et al. 2023. Global Brassicaceae phylogeny based on filtering of 1,000-gene dataset. Current biology: CB 33: 4052–4068.e6.

Hornslien KS, Miller JR, Grini PE. 2019. Regulation of Parent-of-Origin Allelic Expression in the Endosperm. Plant physiology 180: 1498 LP – 1519.

Ishikawa R, Ohnishi T, Kinoshita Y, Eiguchi M, Kurata N, Kinoshita T. 2011. Rice interspecies hybrids show precocious or delayed developmental transitions in the endosperm without change to the rate of syncytial nuclear division. The Plant journal: for cell and molecular biology 65: 798–806.

Johnston SA, Hanneman RE. 1982. Manipulations of endosperm balance number overcome crossing barriers between diploid Solanum species. Science 217: 446–448.

Johnston SA, den Nijs TPM, Peloquin SJ, Hanneman RE. 1980. The significance of genic balance to endosperm development in interspecific crosses. TAG. Theoretical and applied genetics. Theoretische und angewandte Genetik 57: 5–9.

Jordon-Thaden IE, Koch MA. 2008. Species richness and polyploid patterns in the genus Draba (Brassicaceae): a first global perspective. Plant ecology & diversity 1: 255–263.

Kinser TJ, Smith RD, Lawrence AH, Cooley AM, Vallejo-Marín M, Conradi Smith GD, Puzey JR. 2021. Endosperm-based incompatibilities in hybrid monkeyflowers. The Plant cell 33: 2235–2257.

Klosinska M, Picard CL, Gehring M. 2016. Conserved imprinting associated with unique epigenetic signatures in the Arabidopsis genus. Nature Plants 2: 16145.

Krassowski M, Arts M, Lagger C, Max. 2022. krassowski/complex-upset: v1.3.5. Zenodo.

Lafon-Placette C, Hatorangan MR, Steige KA, Cornille A, Lascoux M, Slotte T, Köhler C. 2018. Paternally expressed imprinted genes associate with hybridization barriers in Capsella. Nature Plants 4: 352–357.

Lafon-Placette C, Johannessen IM, Hornslien KS, Ali MF, Bjerkan KN, Bramsiepe J, Glöckle BM, Rebernig CA, Brysting AK, Grini PE, et al. 2017. Endosperm-based hybridization barriers explain the pattern of gene flow between Arabidopsis lyrata and Arabidopsis arenosa in Central Europe. Proceedings of the National Academy of Sciences of the United States of America 114: E1027.

Lafon-Placette C, Köhler C. 2014. Embryo and endosperm, partners in seed development. Current opinion in plant biology 17: 64–69.

Lafon-Placette C, Köhler C. 2016. Endosperm-based postzygotic hybridization barriers: developmental mechanisms and evolutionary drivers. Molecular ecology 25: 2620–2629.

Leblanc O, Pointe C, Hernandez M. 2002. Cell cycle progression during endosperm development in Zea mays depends on parental dosage effects. The Plant journal: for cell and molecular biology 32: 1057–1066.

Lin BY. 1984. Ploidy barrier to endosperm development in maize. Genetics 107: 103–115.

Liu Y, Jing X, Zhang H, Xiong J, Qiao Y. 2021. Identification of Imprinted Genes Based on Homology: An Example of Fragaria vesca. Genes 12.

Liu J, Li J, Liu H-F, Fan S-H, Singh S, Zhou X-R, Hu Z-Y, Wang H-Z, Hua W. 2018. Genome-wide screening and analysis of imprinted genes in rapeseed (Brassica napus L.) endosperm. DNA research: an international journal for rapid publication of reports on genes and genomes 25: 629–640.

Love MI, Huber W, Anders S. 2014. Moderated estimation of fold change and dispersion for RNA-seq data with DESeq2. Genome biology 15: 550.

Lysak MA, Koch MA. 2011. Phylogeny, Genome, and Karyotype Evolution of Crucifers (Brassicaceae). In: Schmidt R, Bancroft I, eds. Genetics and Genomics of the Brassicaceae. New York, NY: Springer New York, 1–31.

Marie-Orleach L, Brochmann C, Glémin S. 2022. Mating system and speciation I: Accumulation of genetic incompatibilities in allopatry. PLoS genetics 18: e1010353.

Marie-Orleach L, Glémin S, Brandrud MK, Brysting AK, Gizaw A, Gustafsson ALS, Riesenberg LH, Brochmann C, Birkeland S. 2023. How Does Selfing Affect the Pace and Process of Speciation. Cold Spring Harbor perspectives in biology.

Marks GE. 1966. THE ORIGIN AND SIGNIFICANCE OF INTRASPECIFIC POLYPLOIDY: EXPERIMENTAL EVIDENCE FROM SOLANUM CHACOENSE. Evolution; international journal of organic evolution 20: 552–557.

Mergner J, Frejno M, List M, Papacek M, Chen X, Chaudhary A, Samaras P, Richter S, Shikata H, Messerer M, et al. 2020. Mass-spectrometry-based draft of the Arabidopsis proteome. Nature 579: 409–414.

Mulligan GA. 1974. Cytotaxonomic studies of Draba nivalis and its close allies in Canada and Alaska. Canadian journal of botany. Journal canadien de botanique 52: 1793–1801.

Müntzing A. 1933. Hybrid incompatibility and the origin of polyploidy. Hereditas 18: 33– 55.

Murashige T, Skoog F. 1962. A revised medium for rapid growth and bio assays with tobacco tissue cultures. Physiologia plantarum 15: 473–497.

Nowak MD, Birkeland S, Mandáková T, Roy Choudhury R, Guo X, Gustafsson ALS, Gizaw A, Schrøder-Nielsen A, Fracassetti M, Brysting AK, et al. 2020. The genome of Draba nivalis shows signatures of adaptation to the extreme environmental stresses of the Arctic. Molecular ecology resources n/a.

Oneal E, Willis JH, Franks RG. 2016. Disruption of endosperm development is a major cause of hybrid seed inviability between Mimulus guttatus and Mimulus nudatus. The New phytologist 210: 1107–1120.

Petrén H, Thosteman H, Stift M, Toräng P, Ågren J, Friberg M. 2023. Differences in mating system and predicted parental conflict affect post-pollination reproductive isolation in a flowering plant. Evolution; international journal of organic evolution 77: 1019–1030.

Picard CL, Povilus RA, Williams BP, Gehring M. 2021. Transcriptional and imprinting complexity in Arabidopsis seeds at single-nucleus resolution. Nature plants 7: 730–738.

Pignatta D, Erdmann RM, Scheer E, Picard CL, Bell GW, Gehring M. 2014. Natural epigenetic polymorphisms lead to intraspecific variation in Arabidopsis gene imprinting (D Weigel, Ed.). eLife 3: e03198.

R Core Team. 2023. R: A Language and Environment for Statistical Computing.

Raunsgard A, Opedal ØH, Ekrem RK, Wright J, Bolstad GH, Armbruster WS, Pélabon C. 2018. Intersexual conflict over seed size is stronger in more outcrossed populations of a mixed-mating plant. Proceedings of the National Academy of Sciences of the United States of America 115: 11561–11566.

Raza A, Hafeez MB, Zahra N, Shaukat K, Umbreen S, Tabassum J, Charagh S, Khan RSA, Hasanuzzaman M. 2020. The Plant Family Brassicaceae: Introduction, Biology, And Importance. In: Hasanuzzaman M, ed. The Plant Family Brassicaceae: Biology and Physiological Responses to Environmental Stresses. Singapore: Springer Singapore, 1–43.

Rebernig CA, Lafon-Placette C, Hatorangan MR, Slotte T, Köhler C. 2015. Non-reciprocal interspecies hybridization barriers in the Capsella genus are established in the endosperm. PLoS genetics 11: e1005295.

Rieseberg LH, Willis JH. 2007. Plant speciation. Science 317: 910–914.

Ritchie ME, Phipson B, Wu D, Hu Y, Law CW, Shi W, Smyth GK. 2015. limma powers differential expression analyses for RNA-sequencing and microarray studies. Nucleic acids research 43: e47–e47.

Rong H, Yang W, Zhu H, Jiang B, Jiang J, Wang Y. 2021. Genomic imprinted genes in reciprocal hybrid endosperm of Brassica napus. BMC plant biology 21: 140.

Roth M, Florez-Rueda AM, Städler T. 2019. Differences in effective ploidy drive genome-wide endosperm expression polarization and seed failure in wild tomato hybrids. Genetics 212: 141–152.

RStudio Team. 2023. RStudio: Integrated Development Environment for R.

Scott RJ, Spielman M, Bailey J, Dickinson HG. 1998. Parent-of-origin effects on seed development in Arabidopsis thaliana. Development 125: 3329 LP – 3341.

Sekine D, Ohnishi T, Furuumi H, Ono A, Yamada T, Kurata N, Kinoshita T. 2013. Dissection of two major components of the post[zygotic hybridization barrier in rice endosperm. The Plant journal: for cell and molecular biology 76: 792–799.

Siddique S, Endres S, Atkins JM, Szakasits D, Wieczorek K, Hofmann J, Blaukopf C, Urwin PE, Tenhaken R, Grundler FMW, et al. 2009. Myo-inositol oxygenase genes are involved in the development of syncytia induced by Heterodera schachtii in Arabidopsis roots. The New phytologist 184: 457–472.

Skrede I, Brochmann C, Borgen L, Rieseberg LH. 2008. Genetics of intrinsic postzygotic isolation in a circumpolar plant species, Draba nivalis (Brassicaceae). Evolution; international journal of organic evolution 62: 1840–1851.

Solbrig OT. 1980. Demography and evolution in plant populations. Univ of California Press.

Sukno S, Ruso J, Jan CC, Melero-Vara JM, Fernandez-Martinez JM. 1999. Interspecific hybridization between sunflower and wild perennial Helianthus species via embryo rescue. Euphytica/ Netherlands journal of plant breeding 106: 69–78.

Tonosaki K, Sekine D, Ohnishi T, Ono A, Furuumi H, Kurata N, Kinoshita T. 2018. Overcoming the species hybridization barrier by ploidy manipulation in the genus Oryza. The Plant journal: for cell and molecular biology 93: 534–544.

Veselá A, Hadincová V, Vandvik V, Münzbergová Z. 2021. Maternal effects strengthen interactions of temperature and precipitation, determining seed germination of dominant alpine grass species. American journal of botany 108: 798–810.

Walker BJ, Abeel T, Shea T, Priest M, Abouelliel A, Sakthikumar S, Cuomo CA, Zeng Q, Wortman J, Young SK, et al. 2014. Pilon: An Integrated Tool for Comprehensive Microbial Variant Detection and Genome Assembly Improvement. PloS one 9: e112963.

Wang L, Yuan J, Ma Y, Jiao W, Ye W, Yang D-L, Yi C, Chen ZJ. 2018. Rice interploidy crosses disrupt epigenetic regulation, gene expression, and seed development. Molecular plant 11: 300–314.

Warwick SI, Francis A, Al-Shehbaz IA. 2006. Brassicaceae: Species checklist and database on CD-Rom. Plant systematics and evolution = Entwicklungsgeschichte und Systematik der Pflanzen 259: 249–258.

Welch BL. 1947. The generalisation of student’s problems when several different population variances are involved. Biometrika 34: 28–35.

Wickham H. 2016. ggplot2: Elegant Graphics for Data Analysis.

Wickham H, François R, Henry L, Müller K, Vaughan D. 2023. dplyr: A Grammar of Data Manipulation.

Wieczorek K, Golecki B, Gerdes L, Heinen P, Szakasits D, Durachko DM, Cosgrove DJ, Kreil DP, Puzio PS, Bohlmann H, et al. 2006. Expansins are involved in the formation of nematode-induced syncytia in roots of Arabidopsis thaliana. The Plant journal: for cell and molecular biology 48: 98–112.

Williams EG, White DWR. 1976. Early seed development after crossing of Trifolium ambiguum and T. repens. New Zealand journal of botany 14: 307–314.

Wolff P, Weinhofer I, Seguin J, Roszak P, Beisel C, Donoghue MTA, Spillane C, Nordborg M, Rehmsmeier M, Köhler C. 2011. High-Resolution Analysis of Parent-of-Origin Allelic Expression in the Arabidopsis Endosperm. PLoS genetics 7: e1002126.

Wu T, Hu E, Xu S, Chen M, Guo P, Dai Z, Feng T, Zhou L, Tang W, Zhan LI. 2021. clusterProfiler 4.0: A universal enrichment tool for interpreting omics data. The innovation journal: the public sector innovation journal 2.

Xu W, Dai M, Li F, Liu A. 2014. Genomic imprinting, methylation and parent-of-origin effects in reciprocal hybrid endosperm of castor bean. Nucleic acids research 42: 6987–6998.

Yoshida T, Kawanabe T, Bo Y, Fujimoto R, Kawabe A. 2018. Genome-Wide Analysis of Parent-of-Origin Allelic Expression in Endosperms of Brassicaceae Species, Brassica rapa. Plant & cell physiology 59: 2590–2601.

Yu G. 2023. enrichplot: visualization of functional enrichment result. R package version 1.20. 1.

Zhang M, Li N, He W, Zhang H, Yang W, Liu B. 2016a. Genome-wide screen of genes imprinted in sorghum endosperm, and the roles of allelic differential cytosine methylation. The Plant journal: for cell and molecular biology 85: 424–436.

Zhang H-Y, Luo M, Johnson SD, Zhu X-W, Liu L, Huang F, Liu Y-T, Xu P-Z, Wu X-J. 2016b. Parental genome imbalance causes post-zygotic seed lethality and deregulates imprinting in rice. Rice 9: 1–12.

Zhang M, Xie S, Dong X, Zhao X, Zeng B, Chen J, Li H, Yang W, Zhao H, Wang G, et al. 2014. Genome-wide high resolution parental-specific DNA and histone methylation maps uncover patterns of imprinting regulation in maize. Genome research 24: 167–176.

Zhang M, Zhao H, Xie S, Chen J, Xu Y, Wang K, Zhao H, Guan H, Hu X, Jiao Y, et al. 2011. Extensive, clustered parental imprinting of protein-coding and noncoding RNAs in developing maize endosperm. Proceedings of the National Academy of Sciences of the United States of America 108: 20042–20047.

