## Supplemental Figures 1-10 for "Low parental conflict, no endosperm hybrid barriers, and maternal bias in genomic imprinting in selfing *Draba* species"

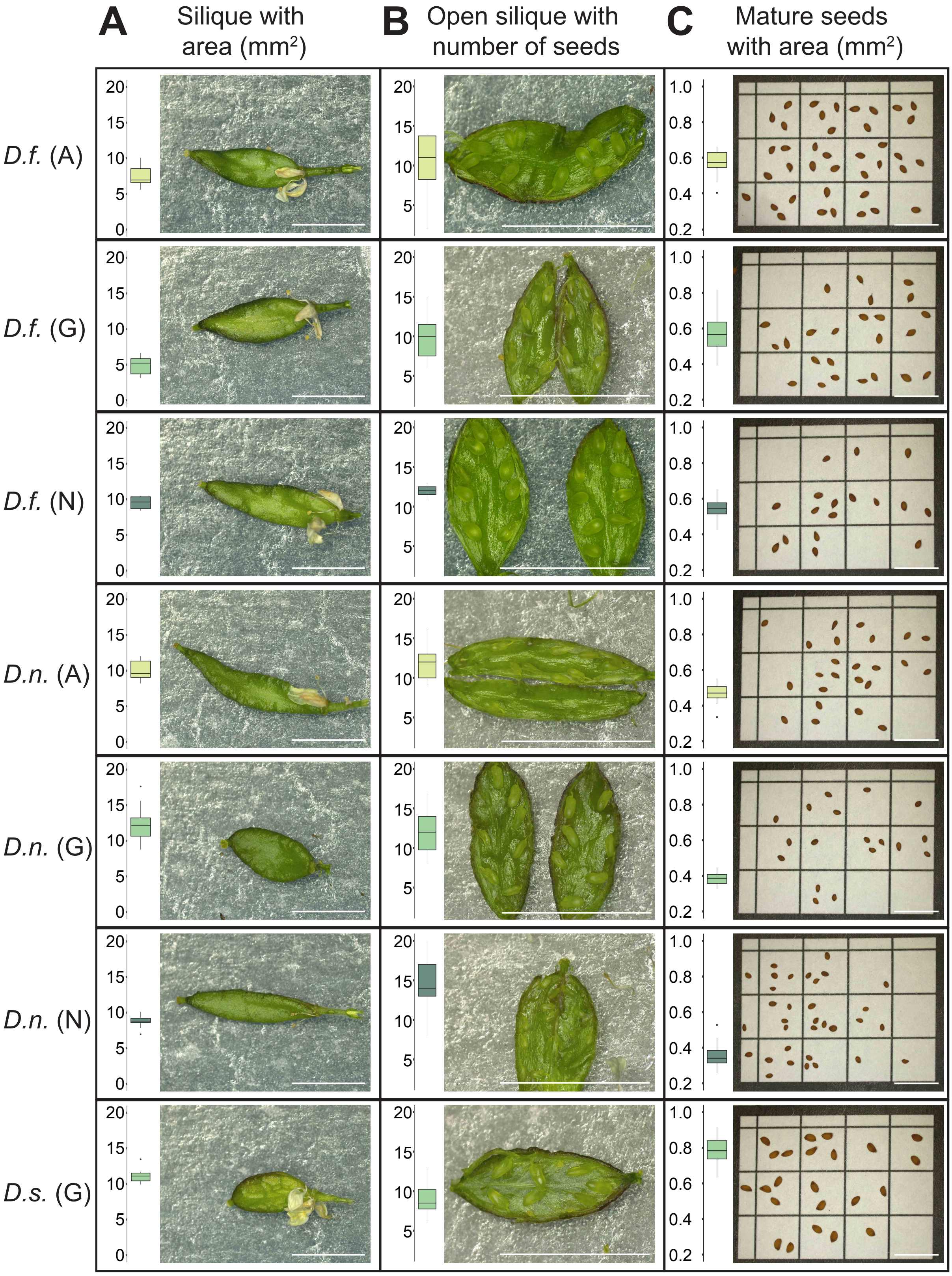


**Supplementary Figure 1: Silique and seed phenotype in *Draba fladnizensis*, *D. nivalis* and *D. subcapitata*. (A)** Stereoscope images of representative siliques with area (mm2) shown by boxplot: *D. fladnizensis* Alaska (*D.f.*(A)), n = 9; *D. fladnizensis* Greenland (*D.f.* (G)), n = 9; *D. fladnizensis* Norway (*D.f.* (N)), n = 9; *D. nivalis* Alaska (*D.n.* (A)), n = 9; *D. nivalis* Greenland (*D.n.* (G)), n = 18; *D. nivalis* Norway (*D.n.* (N)), n = 9 and *D. subcapitata* Greenland (*D.s.* (G)). **(B)** Stereoscope images of immature developing siliques with number of mature seeds per silique shown by boxplot. **(C)** Stereoscope images of mature dry seeds with area (mm2) shown by boxplot: *D.f.* (A), n = 38; *D.f.* (G), n = 23; *D.f.* (N), n = 17; *D.n.* (A), n = 24; *D.n.* (G), n = 18; *D.n.* (N), n = 33 and *D.s.* (G), n = 23. Scale bar = 0.5 cm.

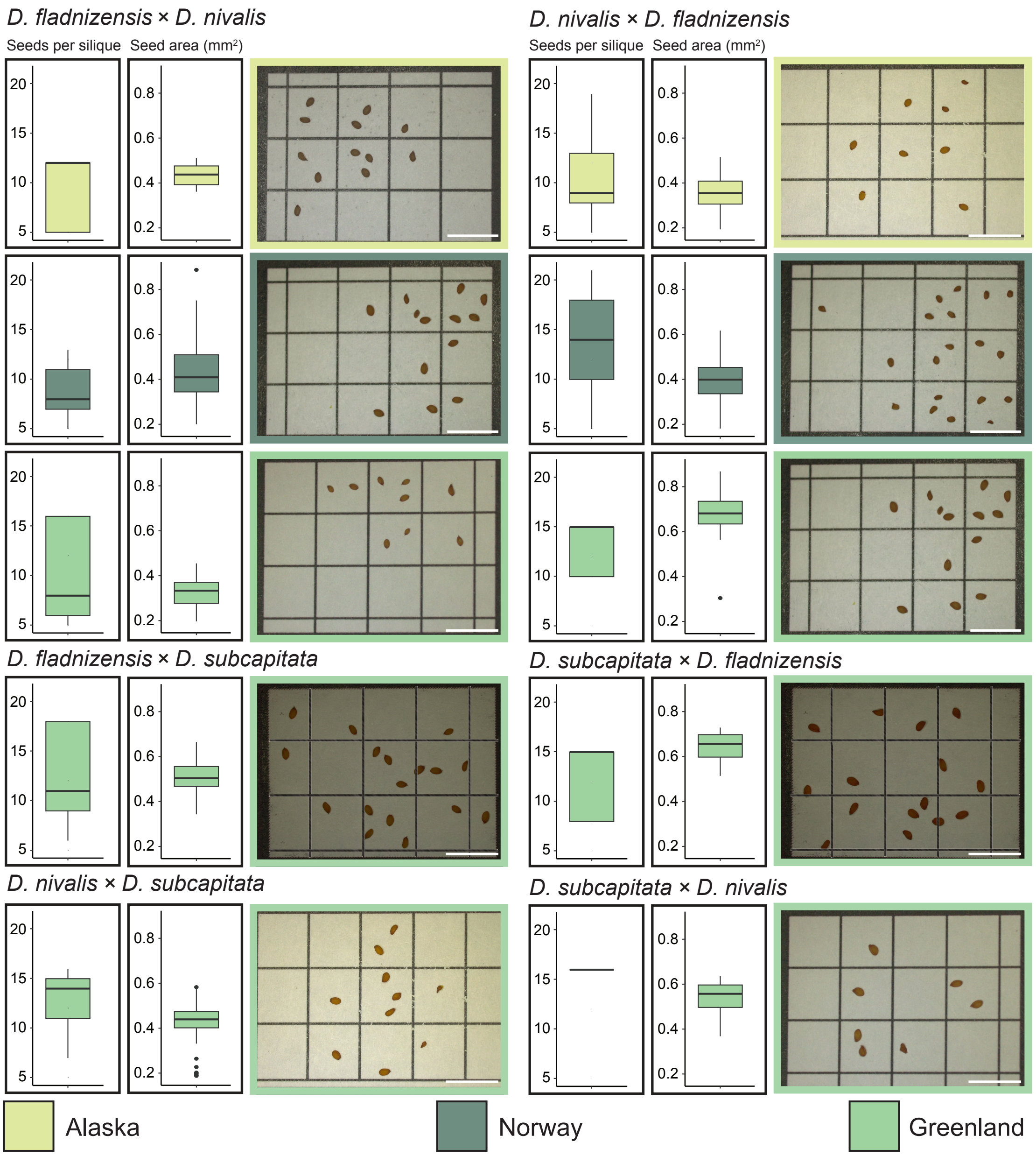


**Supplementary Figure 2: Hybrid seed phenotypes from crosses between *Draba fladnizensis*, *D. nivalis* and *D. subcapitata*.** Seeds per silique, seed area (mm2) and stereoscope images of seeds in: *D. fladnizensis* × *D. nivalis* from Alaska (n = 22), Norway (n = 194) and Greenland (n = 28); *D. nivalis* × *D. fladnizensis* from Alaska (n = 145), Norway (n = 430) and Greenland (n = 25); *D. fladnizensis* × *D. subcapitata* from Greenland (n = 69); *D. subcapitata* × *D. fladnizensis* from Greenland (n = 23); *D. nivalis* × *D. subcapitata* from Greenland (n = 200); *D. subcapitata* × *D. nivalis* from Greenland (n = 7). Outliers are shown as dots. Crosses are indicated as maternal × paternal. Scale bar = 0.5 cm.

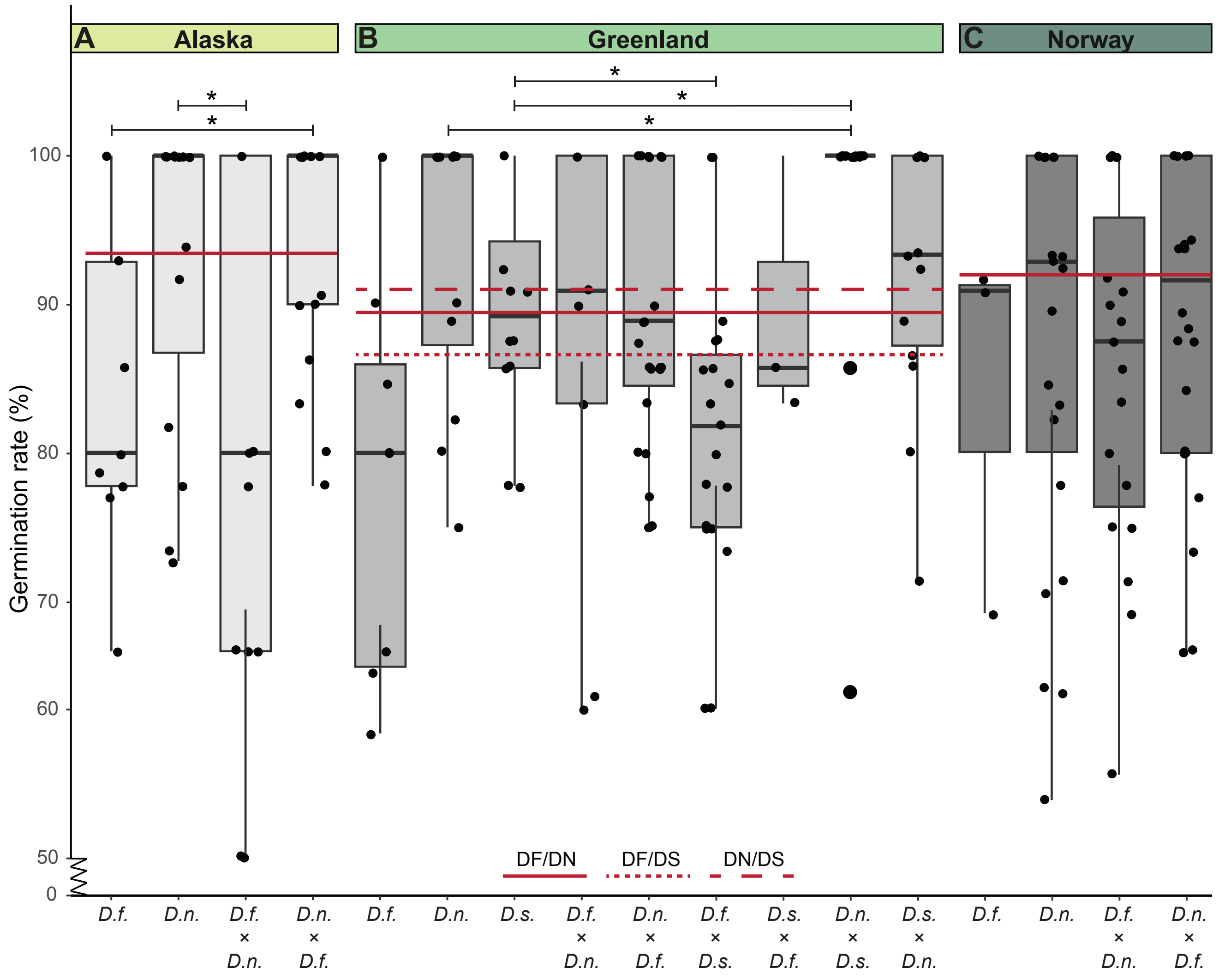


**Supplementary Figure 3: Germination rates of *Draba* hybrids do not indicate endosperm-based hybridization barriers.** Crosses are indicated as maternal × paternal below the x-axis. If the paternal cross partner is not indicated, the maternal and paternal individuals are the same. Biological replicates (n) are given by germination frequency per silique. **(A)** Germination rates of the Alaska populations: *D. fladnizensis* (*D.f.*), n = 9; *D. nivalis* (*D.n.*), n = 15; *D.f.* × *D.n.*, n = 13; *D.n.* × *D.f.*, n = 17. **(B)** Germination rates of the Greenland populations: *D.f.*, n = 8; *D.n.*, n = 12; *D. subcapitata* (*D.s.*), n = 12; *D.f.* × *D.n.*, n = 9; *D.n.* × *D.f.*, n = 27; *D.f.* × *D.s.*, n = 19; *D.s.* × *D.f.*, n = 3; *D.n.* × *D.s.*, n = 17; *D.s.* × *D.n.*, n = 14. **(C)** Germination rates of the Norway populations: *D.f.*, n = 3; *D.n.*, n = 23; *D.f.* × *D.n.*, n = 19; *D.n.* × *D.f.*, n = 24. Biological replicates are plotted as small dots. Outliers are plotted as large dots. Midparent values are indicated as red lines (full, dashed or dotted, depending on the cross, see inset legend above the x-axis). Note that the Y-axis is truncated up to <50%. Significance is indicated for comparisons between reciprocal hybrids and parental lines or midparent value (Welch's t-test: *P ≤ 0.05; not significant is indicated by no asterisk).

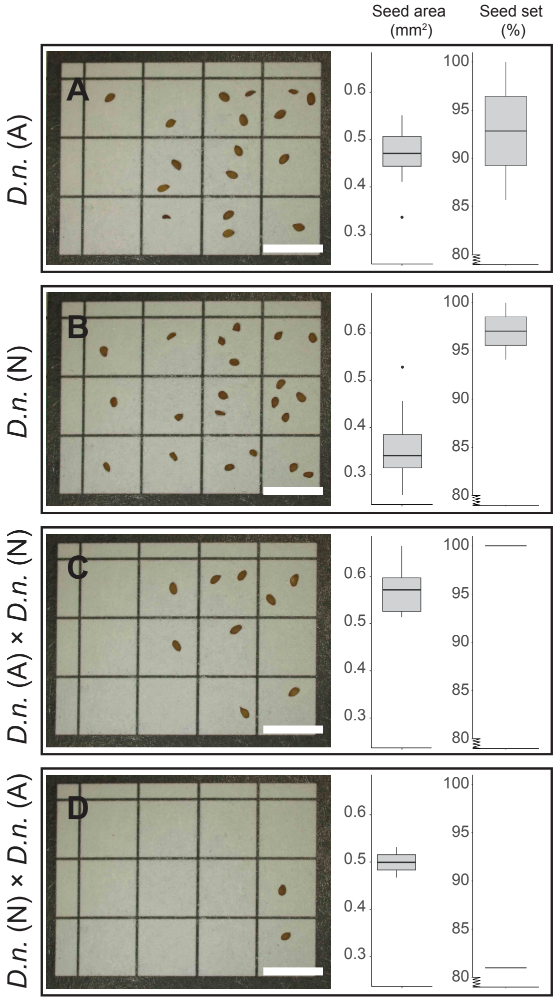


**Supplementary Figure 4: Seed size and seed set of *Draba nivalis* from Alaska and Norway and their reciprocal intraspecific hybrids.** Stereoscope images and seed area (mm2) in: **(A)** *D. nivalis* Alaska (*D.n.* (A)); n = 24, **(B)** *D.n.* Norway (N); n = 33, **(C)** *D.n.* (A) × *D.n.* (N); n = 8 and (D) *D.n.* (N) × *D.n.* (A); n = 2. Seed set shown for the same crosses, based on two siliques for the parental lines and one silique per intraspecific hybrid. Note that the Y-axis is truncated from <80%. Outliers are shown as dots. Crosses are indicated as maternal × paternal. If the paternal cross partner is not indicated, the maternal and paternal individuals are the same. Scale bar = 0.5 cm.

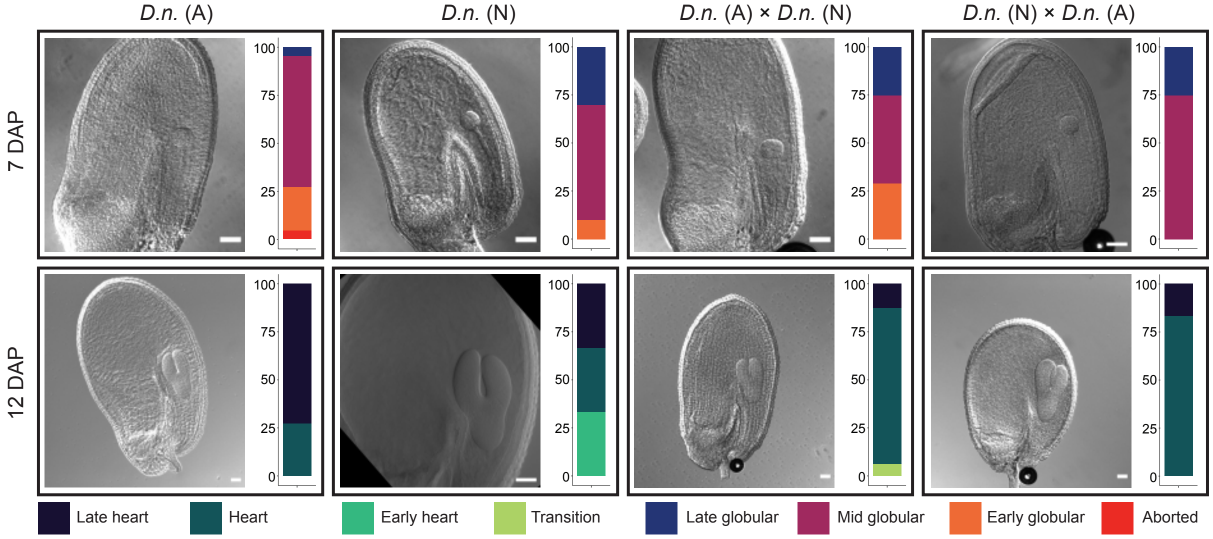


**Supplementary Figure 5: Uniform seed development in *Draba nivalis* from Alaska and Norway.** Light micrographs of bleached seeds at 7 and 12 days after pollination (DAP) of *D. nivalis* Alaska (*D.n.*(A)), *D. nivalis* Norway (*D.n.*(N)), *D.n.*(A) × *D.n.*(N) and *D.n.*(N) × *D.n.*(A). Relative frequencies of embryo stages from the same crosses are shown to the right of each seed, n = 16. Crosses are indicated as maternal × paternal. If the paternal cross partner is not indicated, the maternal and paternal individuals are the same. Scale bar = 50 µm.

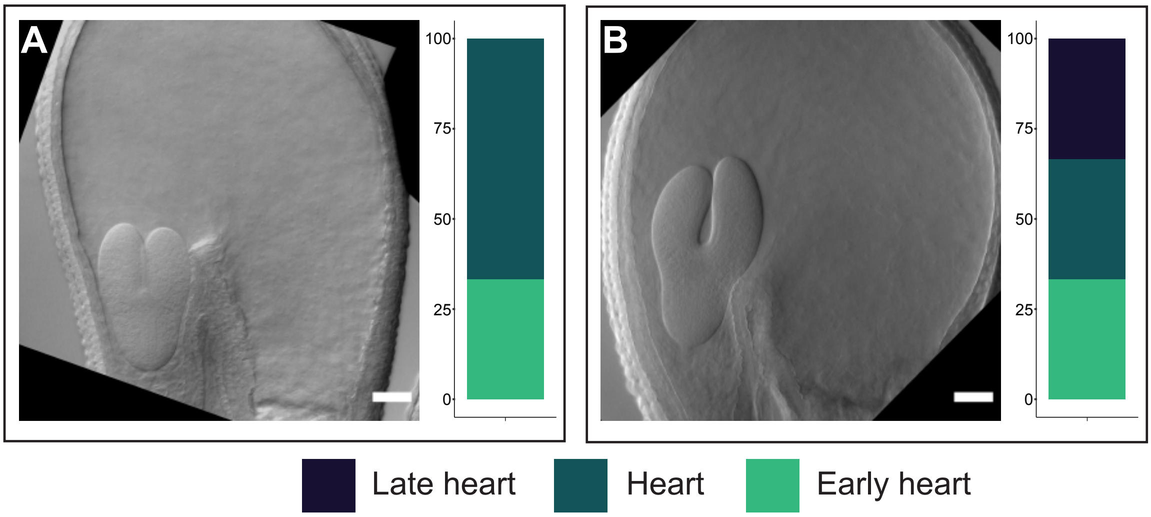


**Supplementary Figure 6: Comparable seed development between the two Norway populations of *Draba nivalis*.** Light micrographs of bleached seeds at 12 days after pollination of **(A)** *D. nivalis* (*D.n.*) Grimsdalen and **(B)** *D.n.* Juvasshytta. Relative frequencies of embryo stages in seeds are shown to the right: *D.n.* Grimsdalen, n = 8; and *D.n.* Juvasshytta, n = 16. Scale bar = 50 µm.

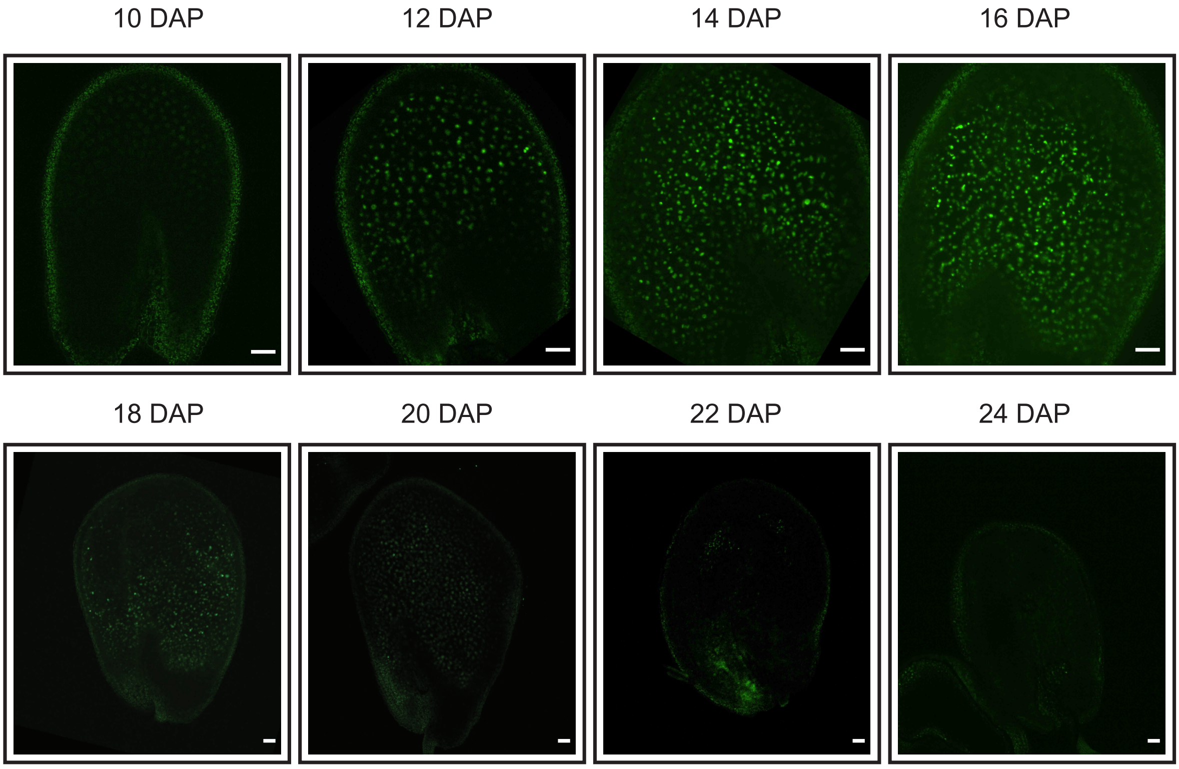


**Supplementary Figure 7: Expression of proAT4G00220>>H2A-GFP in *Draba nivalis* continues until the endosperm is consumed by the embryo.** Confocal micrographs of proAT4G00220>>H2A-GFP (TE1-GFP) in seeds of *Draba nivalis* (N, Grimsdalen) at stages 10, 12, 14, 16, 18, 20, 22 and 24 days after pollination (DAP). Weak GFP expression initiates at 10 DAP, which increases in intensity at 12 DAP and continues until 16 DAP. Thereafter the endosperm is consumed by the embryo and the GFP intensity diminishes. Scale bar = 50 μm.

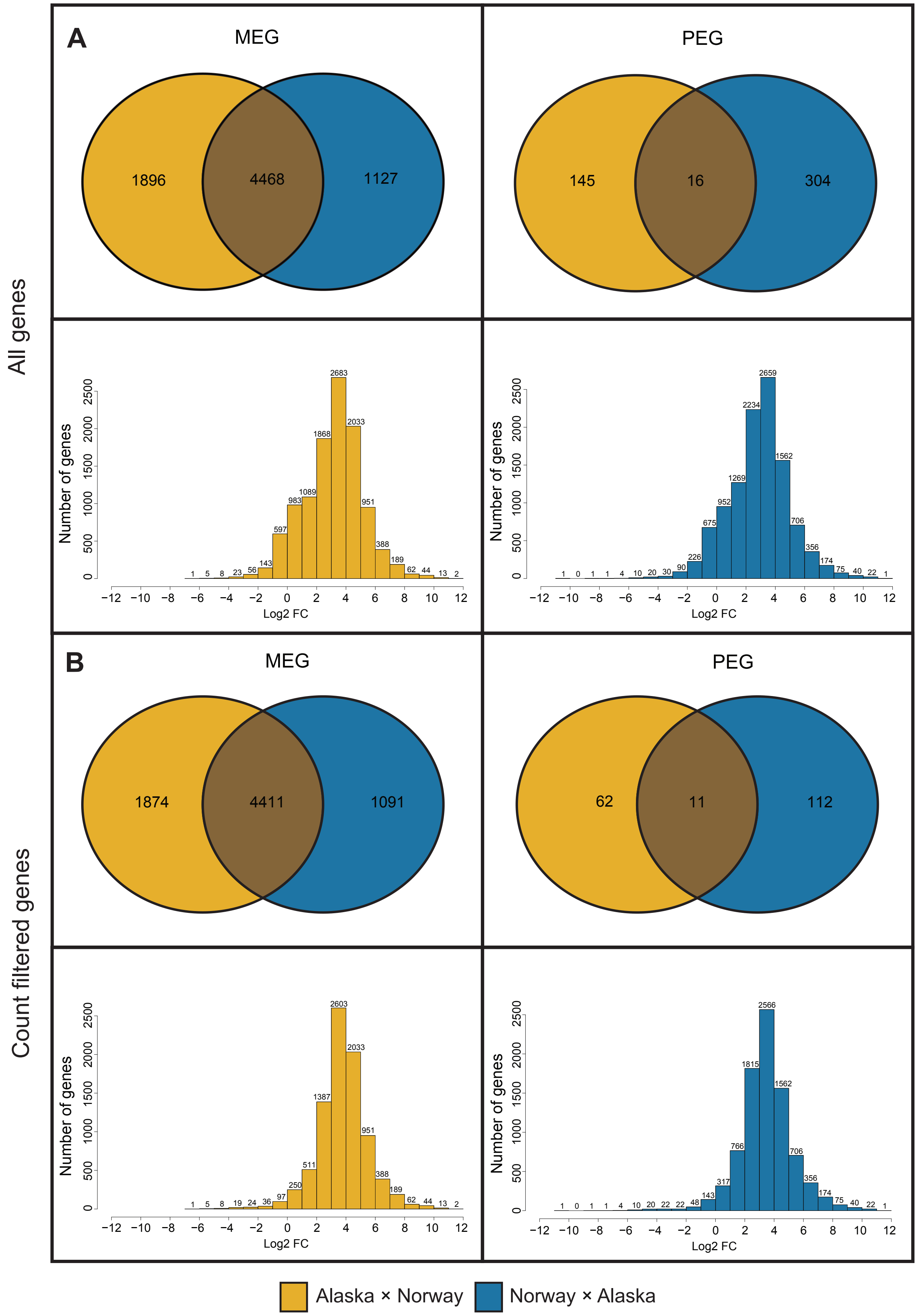


**Supplementary Figure 8: Venn diagrams and histograms of imprinting results in *Draba nivalis* before any filtering and after removal of low-read genes.** Venn diagrams and histograms of maternally expressed genes (MEGs) and paternally expressed genes (PEGs) of the intraspecific *D. nivalis* hybrids: Alaska × Norway and Norway × Alaska, when using all genes with informative reads **(A)**, and after filtering out genes with less than 40 mapped reads, excluding pseudocount **(B)**. Crosses are indicated as maternal × paternal.

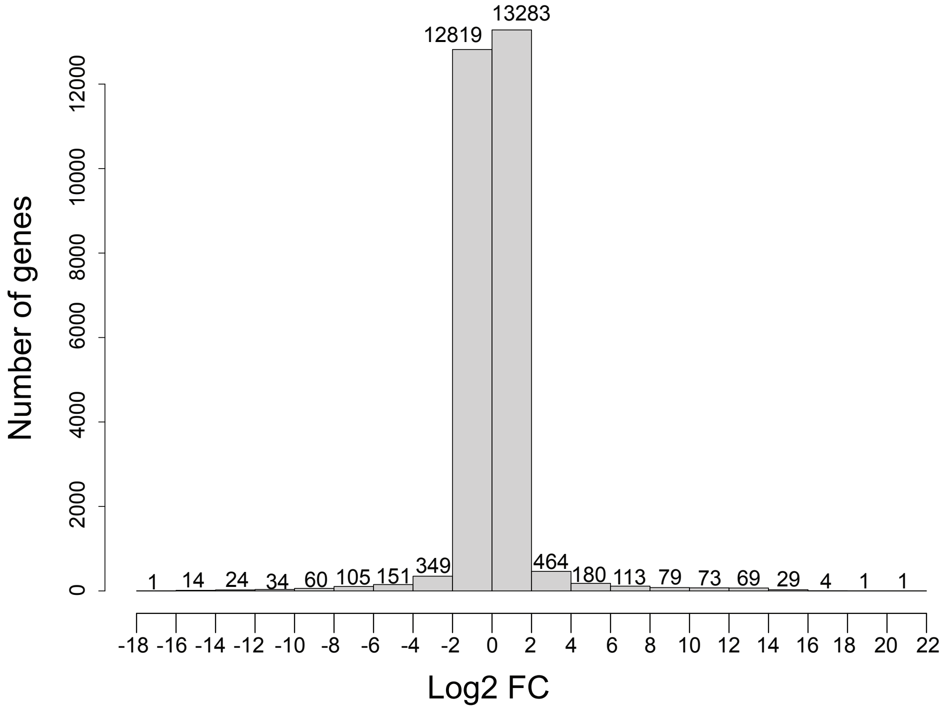


**Supplementary Figure 9: Differentially expressed genes between *Draba nivalis* from Alaska and Norway.** Histogram showing distribution of log2 fold change (FC) values from the differential gene expression analysis (DESeq2) between the two *D. nivalis* populations, Alaska and Norway.

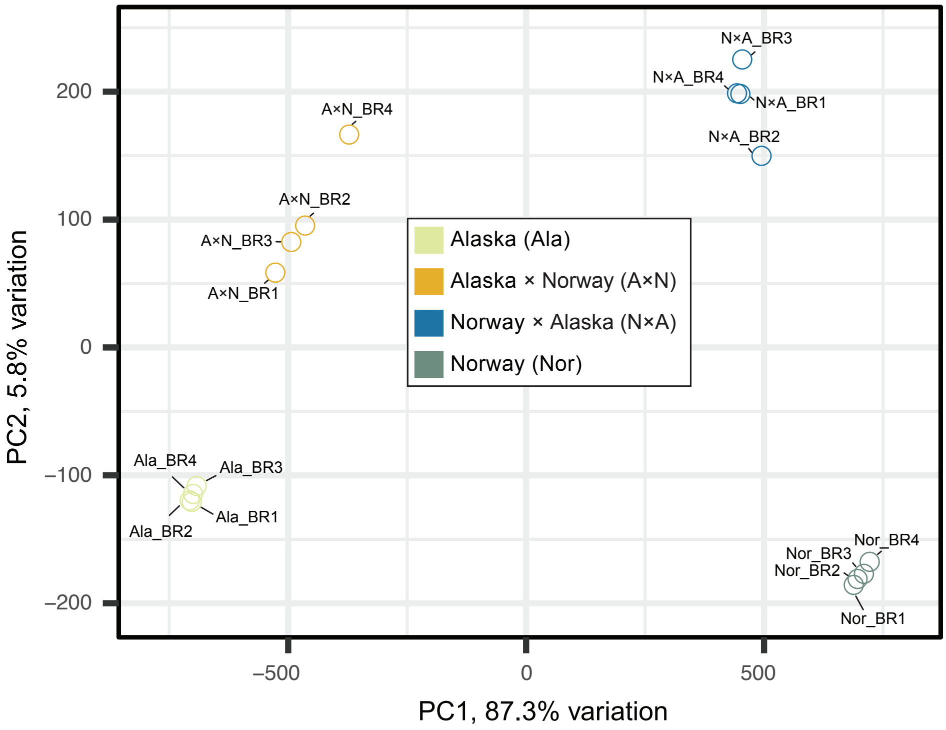


**Supplementary Figure 10: Homogeneity assessment of the Alaska and Norway populations of *Draba nivalis* where general seed coat genes are removed.** Principal component analysis (PCA) based on gene expression of individual biological replicates from the two parental lines of *D. nivalis* from Alaska and Norway and their intraspecific hybrids. Crosses are indicated as maternal × paternal. Replicates from parental lines and intraspecific hybrids align homogeneously.
